# Highly clustered *de novo* frameshift variants in the neuronal splicing factor *NOVA2* result in a specific abnormal C terminal part and cause a severe form of intellectual disability with autistic features

**DOI:** 10.1101/858696

**Authors:** Francesca Mattioli, Gaelle Hayot, Nathalie Drouot, Bertrand Isidor, Jérémie Courraud, Frederic Tran Mau-Them, Chantal Sellier, Maria-Victoria Hinckelmann, Alica Goldman, Aida Telegrafi, Alicia Boughton, Candace Gamble, Sebastien Moutton, Angélique Quartier, Nolwenn Jean, Paul Van Ness, Sarah Grotto, Sophie Nambot, Ganka Douglas, Yue Cindy Si, Jamel Chelly, Zohra Shad, Elisabeth Kaplan, Richard Dineen, Christelle Golzio, Nicolas Charlet, Mandel Jean-Louis, Piton Amélie

## Abstract

The Neuro-Oncological Ventral Antigen 2 NOVA2 protein is a major factor regulating neuron specific alternative splicing, previously associated with an acquired neurologic condition, the paraneoplastic opsoclonus-myoclonus ataxia (POMA). We report here six individuals with *de novo* frameshift variants in the *NOVA2* gene affected with a severe neurodevelopmental disorder characterized by intellectual disability (ID), motor and speech delay, autistic features, hypotonia, feeding difficulties, spasticity or ataxic gait and abnormal brain MRI. The six variants lead to the same reading frame, adding a common 133 aa long proline rich C-terminus part instead of the last KH RNA binding domain. We detected forty-one genes differentially spliced after NOVA2 inactivation in human neural cells. The mutant NOVA2 protein shows decreased ability to bind a target RNA, to regulate specific splicing events and to rescue the phenotype of altered retinotectal axonal pathfinding induced by loss of NOVA2 ortholog in zebrafish. Our results suggest a partial loss-of-function mechanism rather than a full heterozygous loss of function, although a specific contribution of the novel C terminal extension cannot be excluded on the basis of the genetic findings.

## INTRODUCTION

Many genes that play important roles in the development of the nervous system undergo alternative splicing (AS) to generate protein variants with different functions. The use of alternative mRNA isoforms is a critical process crucial during neuronal development, since specific proteins are required at different time and space. Such mRNA diversity is coordinated by different RNA-binding proteins (RBPs). NOVA (Neuro-Oncological Ventral Antigen) proteins, NOVA1 and NOVA2, are two RBPs involved in neuronal-specific alternative splicing^1–3^. They have been first described as antigens in patients with a paraneoplastic neurologic syndrome (POMA), a acquired autoimune neurological disorder characterized by ataxia with or without opsoclonus-myoclonus, with or without dementia, encephalopathy and cortical deficits along other symptoms^4,5^. The two proteins share three similar KH-domains through which they bind directly to YCAY motifs (where Y stands for a pyrimidine) in the messenger RNA (mRNA) sequence^6–8^. According to the binding location on mRNA, they can either induce exon skipping or exon retention ^9^.

Both NOVA1 and NOVA2 are mainly expressed in the central nervous system. However, they show differential expression in specific brain regions in mouse^4^. For instance, NOVA2 is primarily expressed in cortex and hippocampus, whereas NOVA1 is mainly present in midbrain and spinal cord ^10,11^. Both the *Nova1* and the *Nova2* null mice manifest growth retardation, progressive motor dysfunction and death shortly after birth while corpus callosum agenesis is present only in *Nova2*^−/−^ mice ^2,11^. This peculiar defect suggested that NOVA1 and NOVA2 are controlling different set of RNA transcripts. As a matter of fact, NOVA2 seems to be mainly associated with a splicing regulation of genes involved in axonal guidance and projection during the development of a mouse cortex (E18.5), as well as genes implicated in cerebellar function or synapse formation ^11,12^. These alternative splicing (AS) events are developmentally regulated between E12.5 and E18.5 in mouse cortex, highlighting an important role of NOVA2 as an axonal pathfinder modifier during cortical development.

Intellectual disability (ID) is a group of neurodevelopmental disorders (NDD) characterized by significant limitations in both intellectual functioning and adaptive behaviour reflected by an intellectual quotient below 70 before the age of 18. It is genetic in origin in a majority of the cases and these genetic anomalies include chromosomal abnormalities, copy number variants, and point mutations or small insertions/deletions (indels) affecting one of the thousands of genes identified so far as causing ID/NDD when mutated^13^. Large-scale sequencing studies combined with international data exchange allow identification of many novel genes. We report here six individuals with frameshift variants in *NOVA2* that cluster between sequences encoding the second and the third KH domains. We showed that NOVA2 regulates a series of AS events linked to neurite outgrowth and axonal projections in human neural stem cells (hNScs), and that mutant proteins lose, at least partially, the ability to regulate these splicing events. Finally, we demonstrated that inactivation of NOVA2 ortholog impaired axon outgrowth *in vivo* using a zebrafish model and that mutant proteins, on the contrary to wild-type NOVA2, failed to rescue this phenotype.

## RESULTS

### Identification of *de novo* frameshift variants in *NOVA2* leading to expressed truncated proteins sharing the same C-terminal part

We identified a *de novo* frameshift variant, c.782del p.(Val261Glyfs135*), in the *NOVA2* gene (MIM601991) in a an individual with severe intellectual disability (ID) (**Individual 1)**. We collected five additional unrelated individuals with ID with indel variants in *NOVA2*, identified by exome sequencing (ES) by different molecular diagnostic laboratories or research centers (Figure 1a-c). The compilation of these variants resulted from an international collaborative effort partly facilitated by the web-based tool GeneMatcher ^14^. All individuals were initially and *a posteriori* clinically assessed by at least one expert clinical geneticist. **Individual 2** carries a c.711_712insTGT>G p.(Leu238Cysfs*159) found by proband-only WES. Two other deletion/insertion variants causing frameshift in *NOVA2*, c.701_720dup20 p.(Ala241Profs162*) and c.709_748del40 p.(Val237Profs*146), were respectively identified in **Individual 3** and **4**, through trio-based WES. **Individual 5** was identified with a deletion of a nucleotide adjacent to the first variant identified: c.781del, p.(Val261Trpfs*135). The variant was absent from mother’s DNA, but father’s DNA was not available for testing. **Individual 6** carries a *de novo* insertion c.720_721insCCGCGGATGTGCTTCCAGCC, which leads to a truncated protein p.(Ala241Profs162) with one amino acid difference (Phe245 instead of Cys) with those identified in Individual 3. All these variants were unique events never reported in any public variant databases (dbSNP138, 1000 genome, NHLBI GO Exome Sequencing Project, ExAC, GnomAD). No truncating variants were reported in the general population (GnomAD) in the canonical transcript NM_002516.3, suggesting that *NOVA2* is a gene highly intolerant to loss-of-function variants (0 observed vs 12.3 expected, pLI=0.98). There are no known deletion encompassing *NOVA2* in the general popuation from the Database of Genomic Variants (DGV). However, several large deletions encompassing *NOVA2* together with several dozens of other genes are reported in Decipher and ClinVar database in individuals with ID and/or other developmental defects.

**Figure 1.**
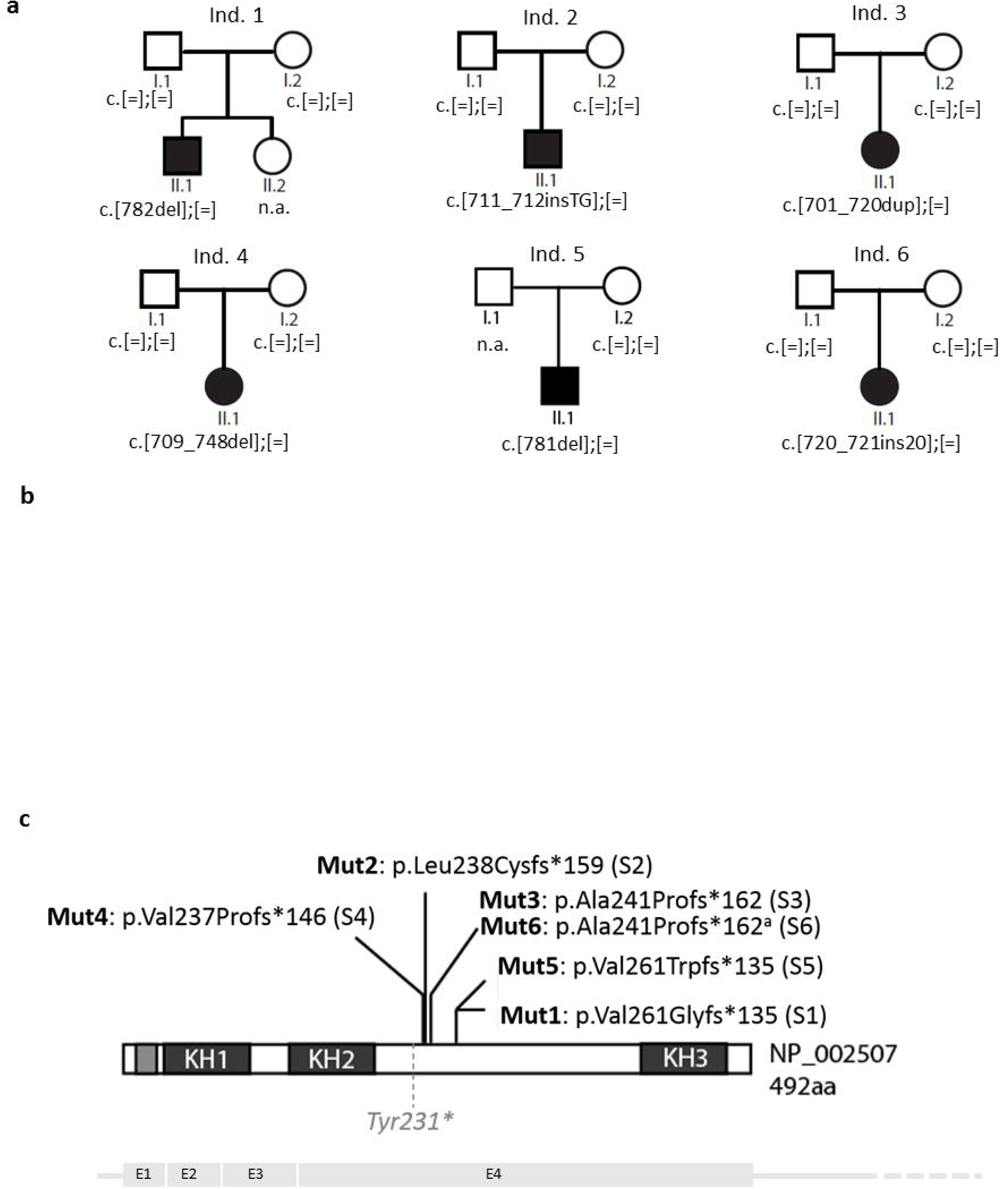
Frameshift variants in *NOVA2* identified in individuals with ID. **(a)** Pedigrees of individuals with frameshift variants in *NOVA2* **(b)** Pictures of Individuals 1,2 and 6. (**c)** Schematic representation of NOVA2 protein showing consequences of the frameshift variants identified in the six patients reported here. Tyr231* indicates a non-existing truncating variant we made for the need of the study. The four coding exons (E1-4) are represented below the protein.

### Frameshift variants in *NOVA2* lead to expressed truncated proteins sharing the same C-terminal part

All six variants are located in the last and largest exon of the gene and lead to frameshifts starting from the same region of the protein (between amino acids 237 and 261). They are predicted to remove the third KH domain of the protein (KH3, amino acids 406-473), which binds RNA loops composed of the tetranucleotide YCAY^7^. The location of all the variants in the last exon of the gene suggested that the mutant transcripts would escape to nonsense-mediated decay (NMD), but the absence of expression in blood prevents confirmation experiment. All the frameshift variants lead to the same alternative frame and the resulting truncated proteins share a common sequence of 133 novel amino acids (**Figure S1-S2**). When overexpressed from plasmid in HeLa cells, mutant NOVA2 protein carrying the variant p.Val261Glyfs*135 identified in Individual 1 (Mut1) is normally expressed and localized (**Figure S3**) with no difference in stability compared to the wild-type protein (cycloheximide treatment, data not shown).

### *NOVA2* inactivation alters axonal guidance *in vivo*

We inactivated *nova1a*, the zebrafish orthologous gene for *NOVA2* (Uniprot ID: Q1LYC7) sharing 78% of sequence identity, using a morpholino (MO). A reduction of the number of axonal tracts formed between optic tecta was observed, as well as a reduction of the tecta size. We counted a total of 29 inter-tecta axonal tracts in control larva whereas the number of inter-tecta axonal tracts was reduced to 10 with an overall altered brain architecture in morphant larva (Figure 2a-b). In vivo complementation assay was performed and we observed that 25pg of human WT *NOVA2* mRNA, but not 25pg of the human Mut1 *NOVA2* mRNA, rescued the MO phenotype. The injection of either the WT or Mut1 *NOVA2* mRNA alone did not affect significantly the number of inter-tecta axonal tracts compared to controls.

**Figure 2.**
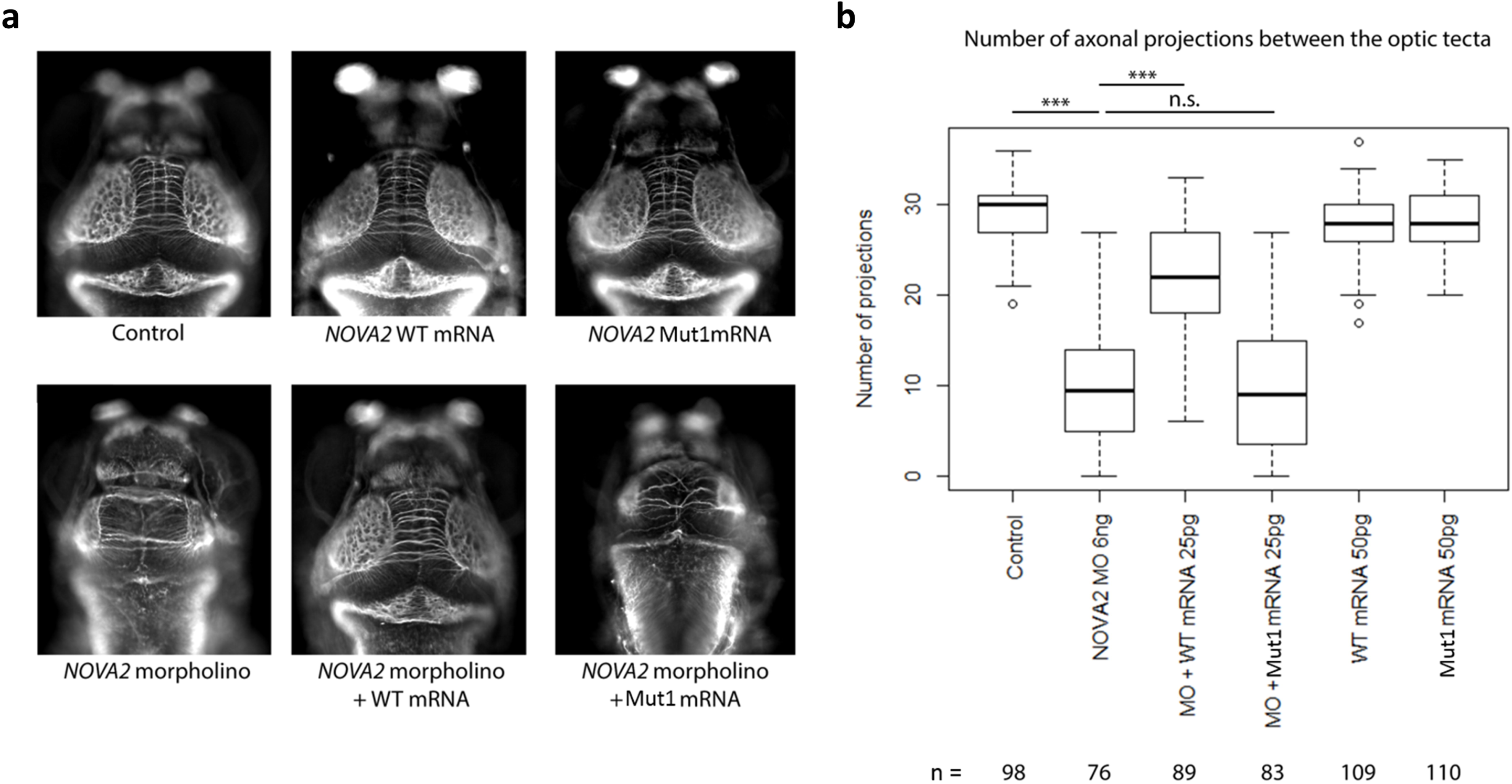
*NOVA2* Mut1 variant leads to decreased number of inter-tecta axonal tracts in zebrafish brain. **(a)** Top panel: Representative images of dorsal views of control, wild-type and Mut1 *NOVA2* mRNA-injected larva at 4 days post-fertilization stained with anti-acetylated tubulin (AcTub). Bottom panel: In vivo complementation assay. Representative images of dorsal views of morpholino (MO)-injected, MO+WT RNA-injected and MO+Mut1 RNA-injected larva at 4 days post-fertilization stained with anti-acetylated tubulin (AcTub) **(b)** Boxplots of inter-tecta axonal tracts’ count after acetylated Tubulin staining of 4 dpf control larva and larva injected with 6ng of *nova1* morpholino (MO), 6ng of *nova1* MO+25pg of WT *NOVA2* mRNA, 6ng of *nova1* MO+25pg of Mut1 *NOVA2* mRNA, 50pg of WT *NOVA2* mRNA, 50pg of Mut1 *NOVA2* mRNA. A t-test was performed between pairs of conditions. p-value < 0.001 are indicated by ***. n.s.: non-significant. n: number of larvae per condition.

### Individuals carrying *de novo* frameshift variants in *NOVA2* present with ID, autistic features and other Angelman syndrome-like clinical manifestations

The main clinical features of the six individuals are summarized in Table 1. More detailed clinical information for all subjects is provided in the Supplemental data. All subjects from the case series exhibited developmental delay (DD) including ID, motor and speech delay, autistic features with stereotypic hands movements and frequent laughter. Other frequent findings included hypotonia (3/4), feeding difficulties (5/6), spasticity or ataxic gait (4/6). Brain imaging showed Chiari malformation type 1 (n=1), cortical atrophy (n=1), and corpus callosum thinning (n=2). Two patients presented with seizures. Notable similarities are found between the reported individuals carrying variants in *NOVA2* and individuals with Angelman syndrome, including feeding difficulties, hypotonia in childhood, inappropriate bouts of laughter, attraction to water, stereotypic movements and severe delay or absence of speech. Interestingly, Angelman syndrome (MIM: 105830) diagnosis was clinically evoked for at least four of the patients, as evident by the prior targeted investigations of the *UBE3A* region by methylation and Sanger sequencing.

**Table 1:**
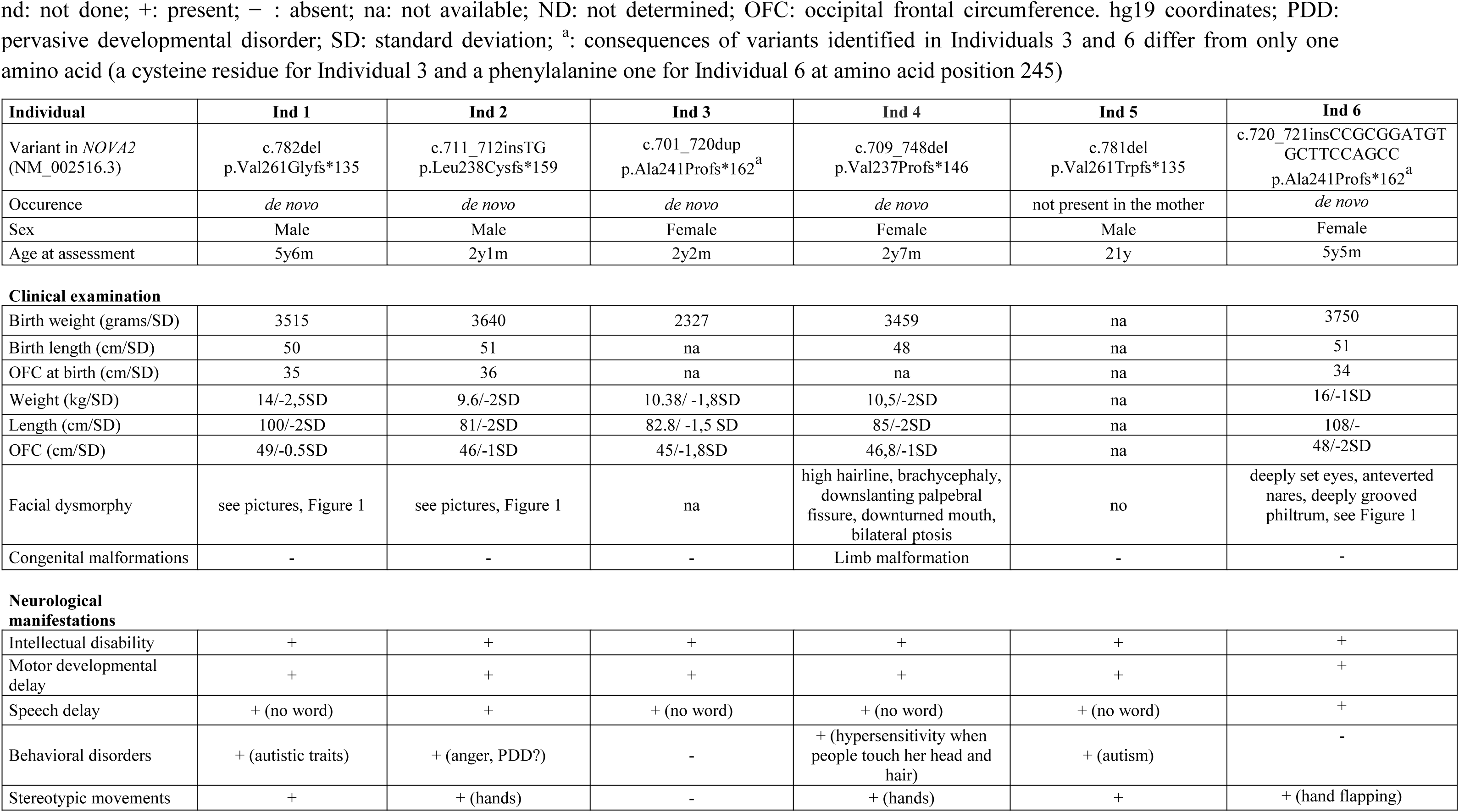
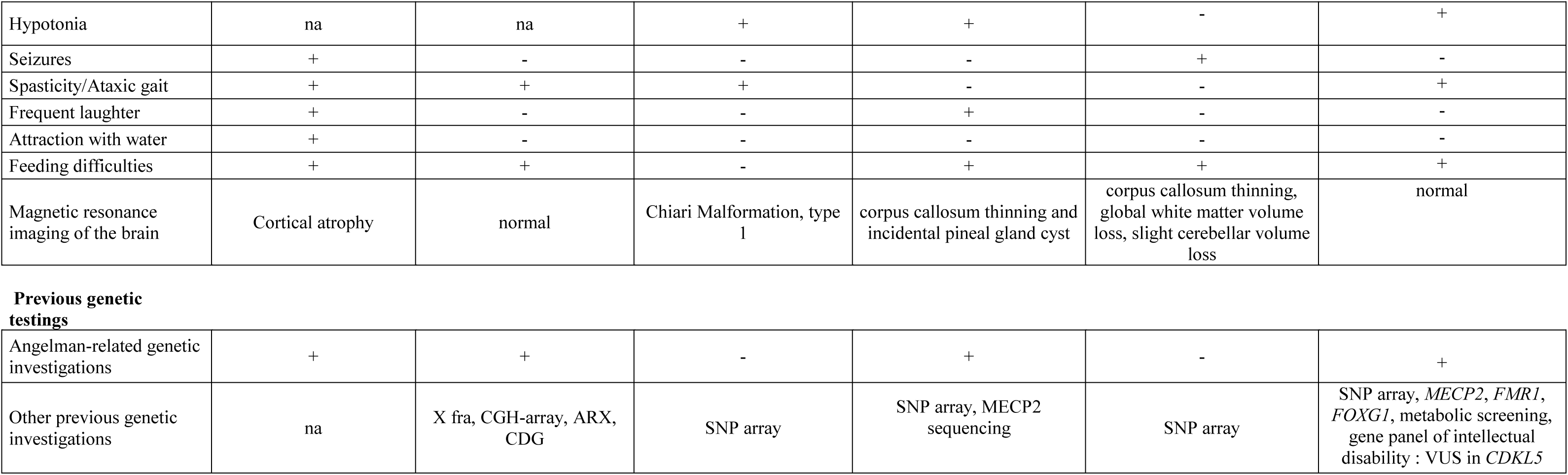
Clinical features of the subjects with frameshift variants in *NOVA2*. nd: not done; +: present; ̶: absent; na: not available; ND: not determined; OFC: occipital frontal circumference. hg19 coordinates; PDD: pervasive developmental disorder; SD: standard deviation; ^a^: consequences of variants identified in Individuals 3 and 6 differ from only one amino acid (a cysteine residue for Individual 3 and a phenylalanine one for Individual 6 at amino acid position 245)

### Identification of splicing events regulated by NOVA2 in human neuronal cells

Most of the data concerning splicing target events regulated by NOVA2 have been generated in mice. In order to identify alternative splicing (AS) events regulated by NOVA2 in human neuronal cells, we used human neural stem cells (hNSCs), self-renewal homogeneous precursors of cortical neurons, derived from embryonic stem cells (SA001). hNSCs were treated with *NOVA2* siRNA, which led to a reduction of 50% in average (+/− 0.09%, n=6) of *NOVA2* mRNA without affecting NOVA1 expression (data not shown). Transcriptomic (RNAseq) analysis revealed that only few genes (<20) were found to be significantly (p-adjusted <0.05) differentially expressed (DE) in hNSCs treated with *NOVA2* siRNA compared to cells treated with the transfection agent only (INTERFERin, INT): 7 up-regulated and 11 down-regulated, with the most significantly one being *NOVA2* (log2 fold change = −0,75, adjusted p-value = 3,62e^−10^)(**Table S1, Figure S4a**). In comparison, no gene was found significantly DE in hNSCs treated with nonspecific siRNA (scramble) compared to INTERFERin (**Figure S4b)**. Splicing analyses with LeafCutter detected AS events significantly different in hNSCs inactivated for *NOVA2* (**Table S2**) in a total of 41 protein-coding genes (Table 2). Enrichment analysis using DAVID (Database for Annotation, Visualization and Integrated Discovery)^15^ demonstrated that the set of genes showing difference in AS events was enriched for Gene Ontology Biological Process terms related to transmembrane proteins and extracellular matrix, cytoskeleton organization, neuron projection/dendrite development (Figure 3a). We found that two third of these genes (26/41) have been previously reported as differently spliced in cortex of *Nova2* Ko mice, including *SGCE*, *NEO1*, *DAB1, SLIT2, SORBS1*, and others^11^. A significant decrease of the skipping of exon 9 from *SGCE* (sarcoglycan epsilon) transcripts was observed for instance in hNSCs after *NOVA2* inactivation (adj. p=1.02e-5, deltaPsy=0.11)(Figure 3b). The effects of *NOVA2* inactivation on the regulation of this AS event were confirmed in another line of hNSCs (GM01869) (Figure 3c). Consistent with these results, we were able to identify several NOVA2 target sequences YCAY in the beginning of *SGCE* intron 9 (Figure 3d). The *SGCE* gene encodes the epsilon-sarcoglycan, a transmembrane protein component of the dystrophin-glycoprotein complex, connecting the actin cytoskeleton to the extracellular matrix, and is known to be submitted to alternative splicing. It causes myoclonus dystonia when mutated ^16^. Among the other AS events significantly affected by *NOVA2* inactivation, we can find for instance a decrease of exon 26 skipping in *NEO1* transcripts (adj. p=0.018, deltaPsy=0.02), which was previously described in mice^17^ (**Figure S5a-b**), a decrease of exon 3 inclusion in *SORBS1* transcripts (adj. p=5e-3, deltaPsy=-0.28)(**Figure S6a-b**) or a decrease of exon 12 skipping for a member of AKAP protein family, *AKAP13* (adj. p=2.6e-4, deltaPsy=0.19) (**Figure S7a-b**). The effects of *NOVA2* inactivation on the regulation of these additional AS events were also confirmed in the GM01869 cell line (**Figure S5c, S6c, S7c**).

**Table 2:**
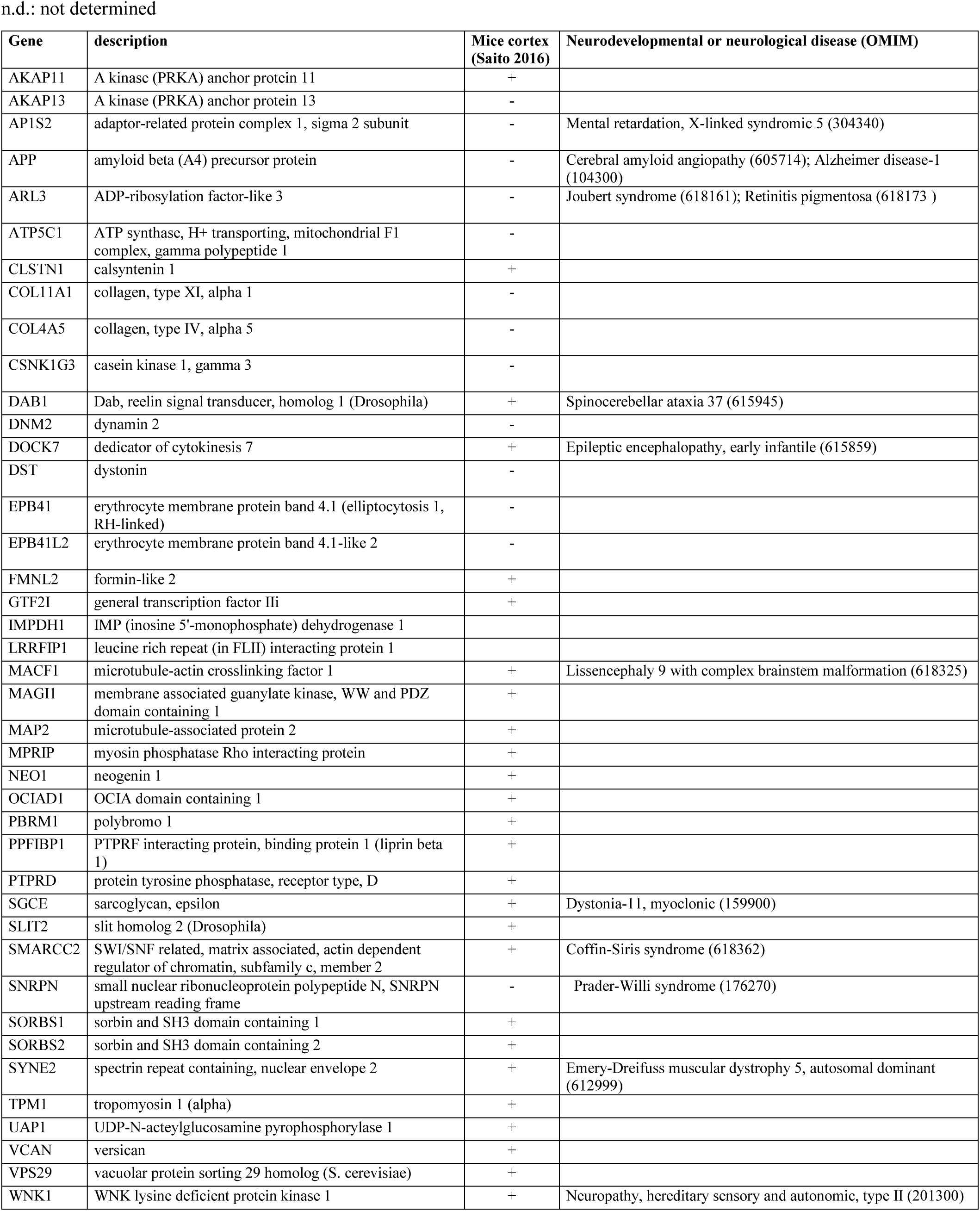
List of genes with alternative splicing (AS) events altered by *NOVA2* inactivation. n.d.: not determined

**Figure 3.**
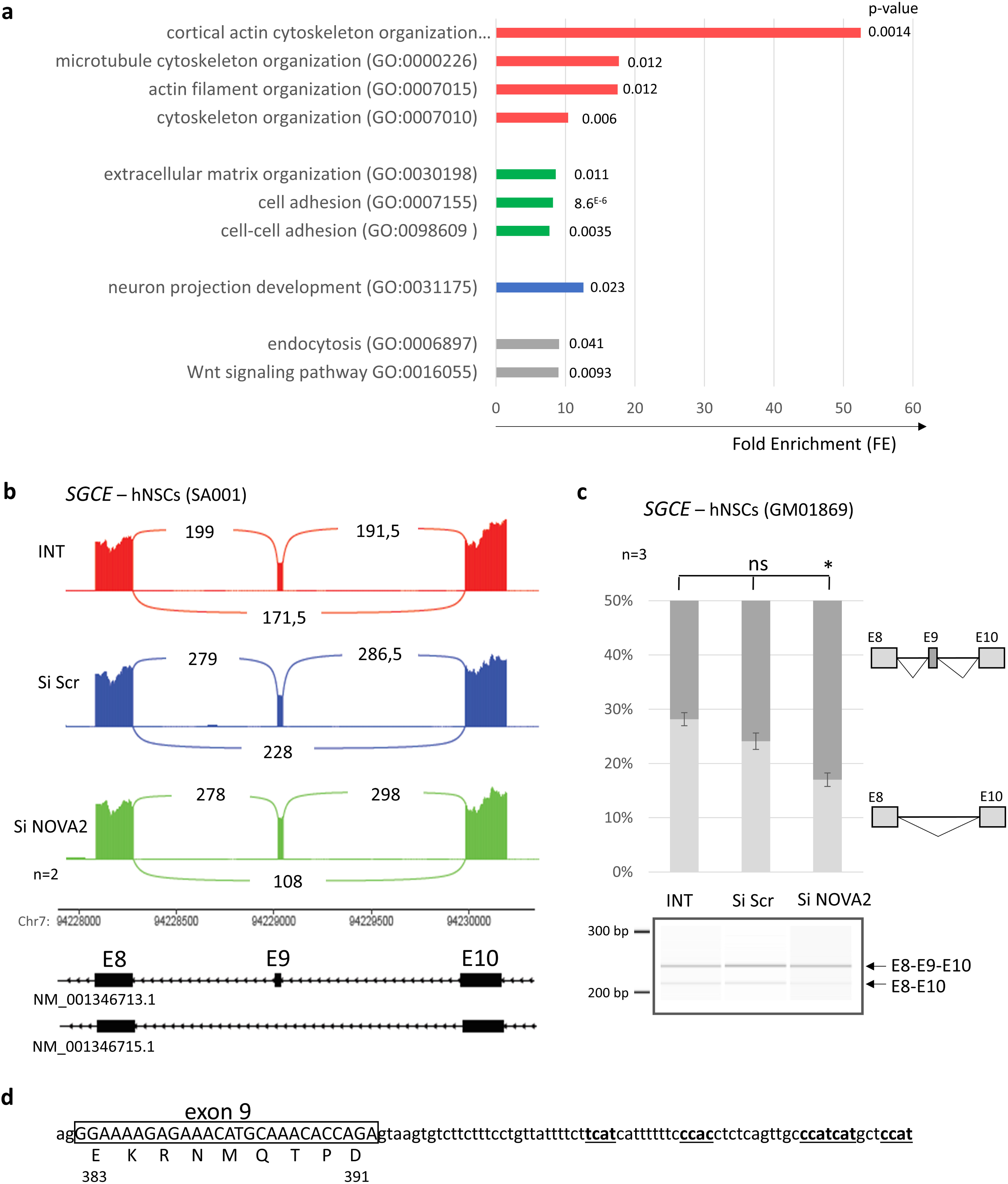
Transcriptomic analysis in human neural stem cells (hNSCs) after *NOVA2* inactivation. **(a)** Enrichment of GO terms (Biological Process and Molecular Function, analyzed using DAVID) for the genes with differential alternative splicing (AS) events identified in human neuronal precursors (hNSCs – SA001) after *NOVA2* inactivation. Fold enrichement (FE) and p-value are also indicated. **(b)** Sashimi plot established from RNA-Seq data representing the AS of *SGCE* exon 9 (NM_001346713.1). The number of reads supporting the existence of each exon-exon junction is indicated as an average between data from the two independent series of hNSCs treated with INTERFERin only (in red), with scramble siRNA (in blue) or with *NOVA2* siRNA (in green) during 48 hours **(c)** Confirmation of the consequences of *NOVA2* inactivation on *SGCE* splicing in another hNSC cell line (GM01869), treated with the transfecting agent only (INT) or transfected with Scramble (si Scr) or *NOVA2* (siNOVA2) siRNA. The RT-PCR products obtained were analyzed by migration on a 2,100 Bioanalyzer instrument (Agilent Technology). Experiments were done in triplicates. The error bars indicate the SEM. Kruskal–Wallis’ ANOVA with Dunn’s multiple comparison test was performed * p<0.05, ns: non-significant. **(d)** Sequence of *SGCE* exon9-intron9 junction with indicated potential NOVA2 binding sequences YCAY.

### Mut1 variant (Val261Glyfs135*) alters NOVA2 ability to bind target sequences and to regulate target alternative splicing (AS) events

Mutant proteins lack the third KH-type domain and we therefore wanted to test their ability to bind RNAs containing the YCAY motif. A gel-shift assays using labeled RNA containing three YCAY sequences showed a significant reduction in RNA binding capacity for NOVA2 Mut1 protein compared to WT NOVA2 (Figure 4a). We then wanted to test the ability of the mutant protein to regulate AS events normally regulated by NOVA2. In HeLa cells, which do not express NOVA2, we observed for instance that most of the *SGCE* transcripts contain exon 9 (Figure 4b). This is consistent with what we observed in hNSCs, where a loss of NOVA2 expression leads to a decrease of exon 9 skipping. The overexpression of WT NOVA2 in HeLa cells leads on the contrary to a significant increase of exon 9 skipping, not observed anymore when Mut1 NOVA2 is overexpressed, demonstrating that the mutant protein has lost its ability to regulate this AS event. We also tested ability of Mut1 NOVA2 to regulate the additional AS events identified in *NEO1*, *SORBS1* or *AKAP13*. The overexpression of WT NOVA2 leads to a significant increase of the skipping of *NEO1* exon 26 and *AKAP13* exon 12, and the inclusion of *SORBS1* exon 5 (**Figure S8**). The complementation assays performed in HeLa cells showed that Mut1 NOVA2 fails to regulate the skipping of *SORBS1* exon 5 but has at least partially preserved activity concerning the regulation of *NEO1* exon 26 and *AKAP13* exon 12 skipping (**Figure S8**). In order to test if the hundred amino acids introduced by the frameshift variants at the C-terminal part of the protein could influence the maintaining of a partial activity of Mut1 NOVA2 protein in the regulation of *NEO1* or *AKAP13* splicing, we introduced in *NOVA2* cDNA a nonsense variant located near to the variants identified in patients, Tyr231*. The resulting truncated protein, which does not contain the hundreds of amino acids added by the frameshift variants, is stably expressed in HeLa cells (**Figure S3**). On the contrary to Mut1 NOVA2, Tyr231* NOVA2 fails to increase the skipping of exon 26 from *NEO1* transcripts and the skipping of exon 12 from *AKAP13* transcripts when it is overexpressed in HeLa cells (**Figure S8**). Coexpression of the WT and Tyr231*, or WT and Mut1 NOVA2 in HeLa cells have a similar effect on AS than the expression of WT alone (**Figure S8**).

**Figure 4.**
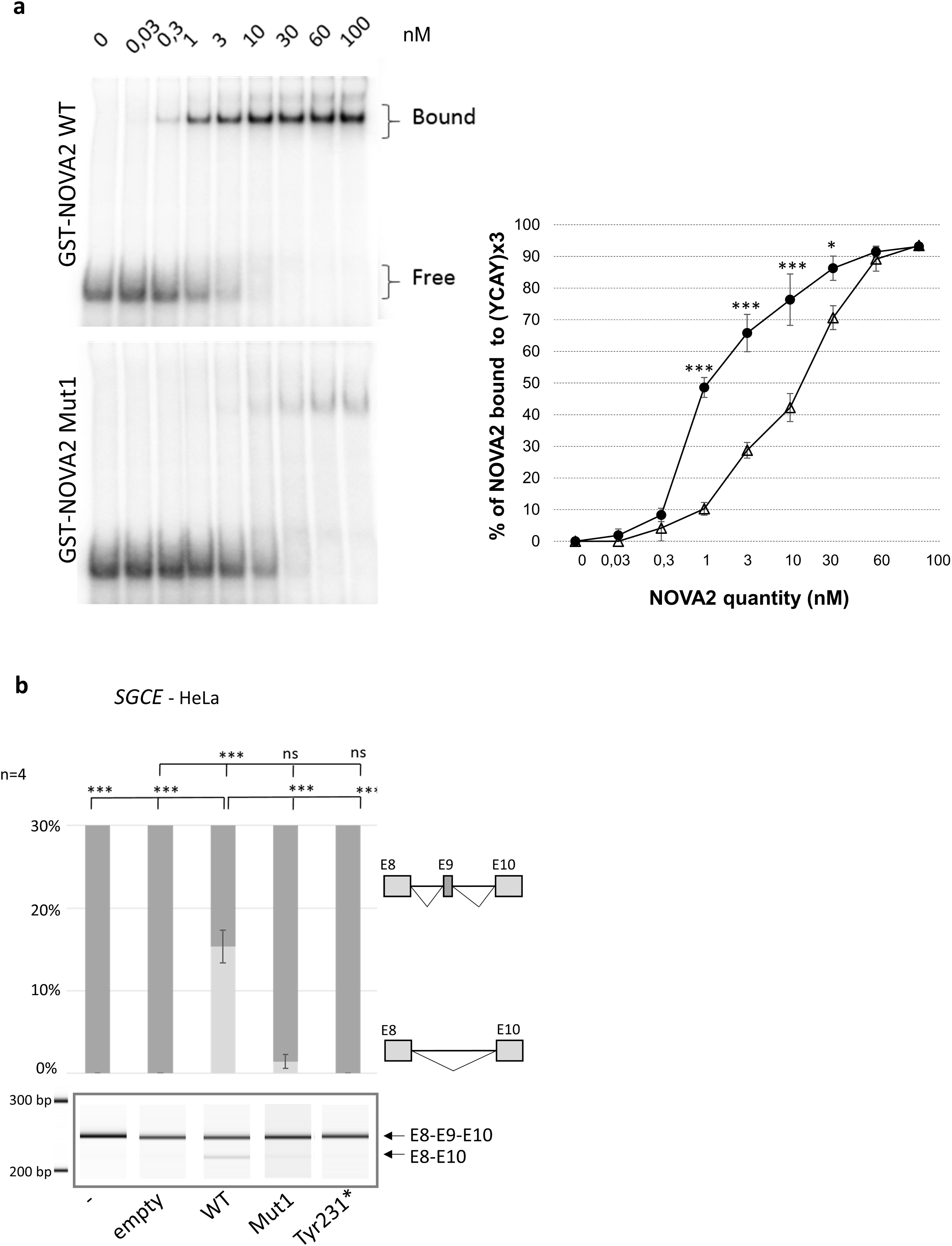
Decreased ability of the Mut1 (Val261Glyfs) NOVA2 protein to bind target RNA sequences (YCAYx3) and to regulate target alternative splicing (AS) events. **(a)** Binding ability of NOVA2 WT and mutant Val261Glyfs (Mut1) proteins to NOVA2-target sequence YCAY 3x. Left panel, gel-shift assays of the indicated amount of purified recombinant GST-NOVA2 WT or Mut1 with 10 pM of uniformly [alphaP^32^] internally labeled *in vitro* transcribed RNAs containing 3 UCAY binding sites for NOVA proteins. Right panel, gel-shift quantification. Error bars standard error mean (SEM) of three independent experiments. Student’s t-test, * indicates p<0.5, *** indicates p<0.001 **(b)** Effect of overexpression of NOVA2 proteins on splicing of SGCE exon 9, in HeLa cells transfected with EGFP-tagged NOVA2 wild-type (WT) or mutant cDNAs. The RT-PCR products obtained were analyzed by migration on a 2,100 Bioanalyzer instrument (Agilent Technology). Four series of experiments were analyzed. The error bars indicate the SEM. Brown-Forsythe and Welch’s ANOVA with Holm-Sidak’s multiple comparisons *** p<0.001, **p<0.01, * p<0.05, ns: non-significant.

## Discussion

We report here a case series of six individuals with severe intellectual disability sharing common autistic features and carrying *de novo* frameshift variants in the same region of the *NOVA2* gene, between two KH RNA-binding domains. This gene encodes an RNA binding protein which is an alternate splicing regulating factor expressed during brain development. We showed that the variant identified in Individual 1 (Mut1 or p.Val261Glyfs135) affects NOVA2 ability to regulate alternative splicing (AS) events and to bind RNA target sequences. All the indel variants cause a shift leading to the same reading frame (frame −1) which results in the addition of a common novel C-terminal part of 133 amino acids from Val261, rich in proline (29.3%) and arginine (12.8%). The alternative frame (frame +1) would have led on the contrary to a more premature stop codon, twenty amino acids downstream of Val261.

To our knowledge, the only case of a specific disorder only caused by such specific distal frameshifts not causing nonsense mediated mRNA decay (NMD) in a single alternate frame resulting in a common long C terminal extension, is the Robinow developmental syndrome (OMIM: 616331 and 616894). This syndrome is characterized by limb and genital anomalies and perturbation of *WNT* signaling, where in the two orthologous genes implicated, the *DLV1* and *DLV3* genes, common proline-arginine rich C terminal extensions of 107 aa (DLV1) or 85 aa (DLV3) have been observed in at least 17 and 6 independent patients, respectively, in the absence of any other type of mutations in these two genes ^18,19^. The authors theorized that the mutant proteins would have a dominant-negative or gain of function effect, although partial loss of function related to the loss of C terminal functional domains were also observed. In the case of indel frameshift variants in *NOVA2*, our functional studies presently support more a loss-of-function effect rather than a dominant negative or gain of function effect. Indeed, no effect was observed when we expressed Mut1 *NOVA2* mRNA alone in zebrafish. On the contrary, Mut1 *NOVA2* mRNA could not rescue the phenotype of loss of axonal projection observed when *NOVA2* is inactivated. The Mut1 protein presents also a partial loss of its splicing regulation activity. Its ability to regulate some AS events is strongly affected in some cases (*SGCE* or *SORBS1)* but only mildly or not affected in others (*NEO1* or *AKAP13*). However, when co-expressed with the WT protein, the mutant NOVA2 does not alter AS, excluding a dominant negative effect of the mutant on AS regulation.

No other LoF variants (frameshift in +1 or nonsense variants) are reported in Clinvar, or in denovo-db database (http://denovo-db.gs.washington.edu/denovo-db/ accessed October 2019). If one takes into account the probability, under a simple heterozygous loss of function hypothesis, that nonsense or canonical splice mutations would also lead to the same phenotype, and under the conservative assumption that frameshifts may be twice more frequent than other types of loss-of-function variants (LoF), the chance of observing only six frameshift variants, all leading to the same frame, and no other LoF variants, is low. We therefore speculate that the longer C-terminal part added by variants leading to frame −1 permits maintenance of a residual activity for NOVA2 protein and that indel variants leading to frame +1 are not observed because they would lead to complete loss of NOVA2 function (with more severe consequences). We corroborated this hypothesis by introducing a stop codon Tyr231* in NOVA2 cDNA: we demonstrated that, if this truncated Tyr231* NOVA2 protein is stably expressed in HeLa cells, it elicits a much stronger loss of AS regulation than for Mut1 NOVA2. However, the consequences of *NOVA2* haploinsufficiency remains puzzling as three individuals carrying deletions encompassing *NOVA2* among several dozens of other genes are reported in the Decipher database. The individuals do not appear to have a more severe phenotype than those reported here, as they present with ID with or without hypotonia, seizures or abnormal gait. Four large deletions (1 to 4 Mb) encompassing many genes in addition to *NOVA2* are also reported in Clinvar. On the other hand it is clear from data in gnomAD that *NOVA2* is a gene with high intolerance to heterozygous LoF in the general population (none observed, pLI = 0.98), and in fact also to missense variants (observed to expected ratio of 0.38 after correction for a deficit in synonymous variants likely due to the poor coverage of part of the gene). Thus we propose that as hypothesized for the Robinow syndrome, the pathomechanism may involve a hypomorphic heterozygous loss-of-function as evidenced by our present results in vitro and in the zebrafish model. However, we could not exclude any gain of function effect linked to the common C-terminal extension that would require longer term experiments in a mouse model to be manifest.

Individuals carrying truncating variants in *NOVA2* exhibited autistic features (stereotypic movements, major speech defect or delay) also overlapping Angelman syndrome (hypotonia, ataxia, seizures and frequent laughter in two patients each), and indeed some of them were prior tested for Angelman syndrome or for Rett syndrome. Four patients presented with anomalies in brain MRI, including corpus callosum thinning in two cases and cortical atrophy. Mouse model inactivated for *Nova2* recapitulate some of these phenotypic observations. Heterozygous KO mouse models present cortical hyperexcitability and epilepsy ^20^ and the homozygous knock-out (KO) mice manifest progressive weakness, motor dysfunction, corpus callosum agenesis, and death shortly after birth^11^. It has recently been reported that conditional mice model with specific inactivation of *Nova2* in Purkinje cells show progressive motor discoordination and cerebellar atrophy^12^.

We demonstrated that NOVA2, like its rodent counterpart, regulates alternative splicing (AS) of different genes encoding proteins involved in signal transduction and cytoskeleton organization, playing a role in neuronal differentiation and migration. In particular, partial inactivation of *NOVA2* leads to mis-regulation of splicing of several transcripts encoding proteins essential for axon outgrowth and pathfinding (*NEO1, SLIT2, etc*). In vivo, the consequences of axonal projection impairments could be illustrated for instance in mammals by corpus callosum hypoplasia (observed in Individuals 4 and 5) or agenesis (previously identified in mice model^21^), but also in zebrafish by the decrease in number of inter-tecta axonal tracts, whose onset involves molecular mechanisms conserved with those of mammals. AS is an essential mechanism aiming at increasing protein diversity and contributes among others to the different critical steps of brain development such as neuronal differentiation, migration, axon guidance, and synaptogenesis. A quarter of the genes identified with AS events mis-regulated when *NOVA2* is partially inactivated are known to cause neurodevelopmental or neurological disorders when mutated, highlighting their crucial role during brain development (Table 2). In addition to *NOVA2*, another gene encoding a protein playing a pivotal role in the regulation of alternative splicing in brain, RBFOX1, also named Ataxin-2-binding protein 1 (A2BP1) or FOX1, is known to be involved in neurodevelopmental disorders. First, its expression was found to be reduced in brains collected postmortem from individuals with ASD leading to alteration of splicing of its predicted targets such as *GRIN1* or *MEF2C*^22^. Moreover, structural variants affecting *RBFOX1* have been reported in several patients with ASD, epilepsy and ID^23–25^.

Mis-regulation of splicing of genes important for brain functioning have also been reported in neurodegenerative diseases, such as Alzheimer disease, fronto-temporal dementia, amyotrophic lateral sclerosis, etc^26^. Interestingly, the targeting of NOVA proteins by an abnormal immune response causes the POMA disease, a neurological condition characterized by ataxia with or without opsoclonus-myoclonus, dementia, encephalopathy and cortical deficits. Other genes encoding RBP have already been implicated in both child neurodevelopmental conditions and adult-onset neurological and/or neurodegenerative disorders. For instance, deletions and missense variants in PUM1 have been reported to cause a global developmental delay syndrome characterized by speech delay, ID, ataxia and seizure, while a rare missense variant with milder functional effect was identified in a family with several members affected by an adult-onset ataxia with incomplete penetrance^27^. FMRP, the gene responsible for the most frequent cause of X-linked ID, the Fragile X syndrome (FXS), has been also implicated in an adult-onset neurodegenerative disease: while large CGG expansion in the 5’UTR of FMR1 cause FXS, expansions with an intermediate number of repeats cause the Fragile-X Tremor Ataxia Syndrome (FXTAS)^28^. Overall, defects in RBPs causing both neurodevelopment and adult-onset neurological/neurodegenerative disorders highlight the importance of post-transcriptional gene expression regulation in neuronal cells for both brain development and brain functioning throughout life.

In conclusion, through a multi-center collaboration, we identified truncating variants in *NOVA2* affecting its activity on AS regulation and responsible for a severe syndromic form of NDD/ID with clinical manifestations overlapping the Angelman syndrome. The frequency of this syndrome is difficult to evaluate. No frameshift variant have been observed in about 10,000 probands with NDD subjected to exome sequencing in the DDD project (Matt Hurles, personal communication) or in 27,000 diagnostic exomes in Netherland (Han Brunner and Rolph Pfundt, personal communication). However, the frameshift variants we report in this study are located in a GC-rich internally repetitive region of NOVA2 that is very poorly represented in most exome sequencing data (**Figure S9**). Moreover, half of the frameshifts we identified are caused by indels of 20 to 40 bp, that are difficult to detect in standard exome protocols, especially when relatively short read lengths are used (75 bp to 100bp). This study highlights the importance of the regulation of AS events during brain development and how alterations of this regulation might lead to brain dysfunction.

## METHODS

### Patients and genetic analyses

**Individual 1** and biological parents were followed at the Service de Génétique Médicale of Nantes Universitary Hospital, and were enrolled for genetic testing in the laboratory of genetic diagnosis of Strasbourg University Hospital. Targeted sequencing of hundred candidate genes for ID did not reveal pathogenic variants ^29^. A family trio-based WES was performed as previously described^30^ using 100bp paired-end sequencing. The NOVA2 *de novo* pathogenic variant c.782del was confirmed by Sanger sequencing and trio compatibility was checked by using polymorphic microsatellite markers (PowerPlex 16HS system, Promega). **Individuals 2 and 6** and biological parents were enrolled for genetic diagnosis in Dijon University Hospital and in Assistance publique – Hôpitaux de Paris (APHP) respectively. Proband only WES was performed as previously described using 75pb paired-end sequencing^31^. **Individuals 3, 4, 5** and their biological parents were enrolled respectively at the following centers; at the Cook Children’s Genetics in Fort Worth, at the Department of Genetics, University of Illinois College of Medicine in Chicago, and at the Department of Neurology, Baylor College of Medicine in Houston, Texas, USA. Trio whole exome sequencing was obtained in a CLIA approved clinical genetic diagnostic laboratory, GeneDx (https://www.genedx.com) as described previously^32^. Sequencing was done on an Illumina system with 100bp paired-end reads. Reads were aligned to human genome build GRCh37/UCSC hg19 and analyzed for sequence variants using a custom-developed analysis tool. The different genetic studies were approved by the local Ethics Committee and written informed consent for genetic testing and authorization for publication were obtained from their legal representative.

### NOVA2 plasmids and siRNA

Human cells were transfected with a pcDNA3.1 plasmid containing the sequence of human *NOVA2* cDNA. The plasmid was optimized to reduce GC content while keeping the amino acid sequence and GFP location at the N-terminal side of the *NOVA2* gene. Mut1 variant (c.782del, p.Val261Glyfs*135), identified in Individual 1, was introduced by site-directed mutagenesis. We also introduced a substitution leading to a premature truncated protein: c.693C>A, Tyr231*. For experiments in zebrafish the human wild-type and mutant (Mut1) *NOVA2* cDNA were introduced into PSC2 plasmids. The sequences of all constructs were confirmed by Sanger sequencing (GATC Biotech, Konstanz, Germany). Pools of siRNA against human *NOVA2* gene and of Scramble siRNA were purchased from Dharmacon (GE Healthcare, Lafayette, CO).

### Cell culture and transfection

Human neuronal stem cells (hNSCs) were derived from human embryonic stem cell line (SA001) and from reprogrammed fibroblasts (GM01869). They were obtained from I-Stem (Cellartis, Goteborg, Sweden, work supervised by the French Bioethics Agency, and Coriell Institute for Medical Research, Camden, NJ) as described previously^33^. hNSCs were seeded on poly-ornithine and laminin-coated dishes and maintained in N2B27 medium (DMEM/F12 and Neurobasal medium (1:1) supplemented with N2, B27, 2-mercaptoethanol (all from Invitrogen, Carlsbad, CA, USA), BDNF (20ng/mL), FGF-2 (10ng/mL) (both from PeproTech, Rocky Hill, NJ, USA) and EGF (R&D Systems; 10ng/mL). Culture and quality controls of hNSCs were performed as described in Quartier et al., 2018^34^. hNSCs were transfected using INTERFER in reverse transfection protocol (Polyplus-transfection, Illkirch-Graffenstaden, France) with scramble siRNA, *NOVA2* siRNA (pool of 20nM each) or transfecting agent only. HeLa cells were maintained in DMEM supplemented by 1g/L glucose with gentamycin and 5% Fetal Calf Serum in a 37°C, 5% CO2 humidified incubator with medium renewed every two days. Cells were transfected at 60-70% of confluence in 6-well plates using Lipofectamine® 2000 DNA transfection reagent (Invitrogen®) in Opti-MEM according to manufacturer’s instructions with 2µg of each *NOVA2* plasmid. Cells were stopped 24/48 hours after transfection for RNA extraction.

### Western blot and immunofluorescence

Four series of HeLa cells were transfected with empty pcDNA3.1 or pcDNA3.1 containing wild-type or mutant (Mut1 and Tyr231*) NOVA2 sequences. To control transfection efficiency, cells were co-transfected using a previously published pcDNA3.1-FLAG-THOC6 constructs.^30^. HeLa cells were lysed in RIPA buffer with protease inhibitor cocktail (Roche) 24h hours after transfection. Proteins were separated after denaturation on a 10% acrylamide gel and transferred onto a PVDF membrane. EGFP-tagged NOVA2 proteins were visualized using an in-house mouse anti-GFP antibody (1:10,000). Their expression levels were normalized using the FLAG staining (FLAG antibody: 1:1,000; F1804, Sigma-Aldrich, Saint-Louis, MO, USA).

### RNA sequencing

Experiments were performed in duplicate from two independent series of hNSCs SA001 treated with the transfecting agent only (INTERFERin) or transfected with scramble siRNA or anti-*NOVA2* siRNA during 48 hours. Total RNA was extracted using the RNeasy mini kit (Qiagen, Valencia, CA, USA) including a DNase treatment levels and quality were quantified using a Nanodrop spectrophotometer and a 2100 Bioanalyzer (Agilent, Santa Clara, CA, USA). RNA mRNA libraries of template molecules suitable for high throughput DNA sequencing were created using the KAPA mRNA HyperPrep Kit (Roche). Briefly, mRNAs were purified from 300ng of total RNA using poly-T oligo-attached magnetic beads and fragmented at 85°C with magnesium for 6 min. The cleaved mRNA fragments were reverse transcribed into cDNA using random primers and KAPA Script enzyme. Second strands were synthesized and adapters were added according to manufacturer’s instructions. Libraries were enriched by PCR amplification (12 cycles). PCR products were purified and size selected (400bp) before sequencing using paired-end 100bp, Illumina Hiseq 4000 sequencer. Sequencing generated on average 100 to 200 million of reads per sample that were mapped onto the hg19 assembly of the human genome using TopHat 2.0.14 ^35^ and the Bowtie 2-2.1.0 aligner^36^. Data are available in Gene Expression Omnibus (GEO), (https://www.ncbi.nlm.nih.gov/geo/: GSE138766). Gene expression was quantified using HTSeq-0.6.1^37^ and gene annotations from Ensembl release 75. Only uniquely-mapped and non-ambiguously assigned reads were retained for further analyses. Read counts were then normalized across libraries with the median-of-ratios method proposed by Anders and Huber ^38^. Relative Log Expression (RLE) plots were drawn to check that the distributions were centered around the zero line and as tight as possible to make sure that normalization was performed correctly. Comparisons to untreated cells (receiving transfection agent, INTERFERin, only) were performed using the statistical method proposed by Anders and Huber (Anders et Huber 2010). The Wald test was used to estimate the p-values and they were adjusted for multiple testing with the Benjamini and Hochberg method ^39^. LeafCutter was used to identify alternative splicing (AS) events between *NOVA2* inactivation and other conditions ^40^. Genes with AS events were analyzed using the Database for Annotation, Visualization and Integrated Discovery (DAVID 6.7). Biological processes and molecular functions of Gene Ontology Consortium (GO) were used for the functional annotations.

### Confirmation of alternative splicing (AS) events identified by RNASeq

GM01869 hNSCs were used to validate results obtained by RNA-Seq in the SA001 line. Cell were treated with tranfecting agent, scramble or *NOVA2* siRNA as described for SA001 (each condition in triplicates). For splicing analyses in HeLa cells, 6-well plates of cells were transfected with 2µg of empty or NOVA2 pcDNA3.1 plasmids alone or combined (three to four series of independent experiments were analyzed). mRNA was extracted 24 hours after siRNA transfection using using the RNeasy mini kit (Qiagen, Valencia, CA, USA). 500ng of total RNA was reverse transcribed into cDNA using random hexamers and SuperScript IV reverse transcriptase according to manufacturer’s recommendation. PCR was performed (16 cycles of touch-down 70°-54° followed by 8 to 21 cycles at 59° depending of the mRNA), using the following primers NEO1_E25_F 5’-GGAAGGCGAGGAATGAGACCAAAA-3’ and NEO1_E27_R 5’-TGTGCTTGGCAATGCAGGATCA-3’, AKAP13_E11_F 5’-AAGTGCCTGCAAACTGCTCTGT-3’ and AKAP13_E13_R 5’-AAAGAGTCAACCCGTTCCTCACCA-3’, SORBS1_E2_F 5’-TGTGATGAATGGCTTGGCAC-3’ and SORBS1_E4_R 5’-TCCCTTCCCAGTGCAGATTT-3’, SGCE_E8_F 5’-TGGTGGAGAATACAAACCCC-3’ and SGCE_E10_R 5’-GGACATGTCTCGAAGCTCCT-3’. The PCR products obtained were analyzed by migration on a 2,100 Bioanalyzer instrument (Agilent Technology).

### Electrophoretic mobility shift assay

In order to produce recombinant NOVA2 proteins, E. coli BL21 (RIL) pRARE competent cells (Invitrogen) were transformed with pet28a-GST-NOVA2 WT or Mut1. The cells were then incubated at 30°C in 400 ml of LB medium supplemented with Kanamycin until an OD600 of 0.5. Afterwards, 0.5 mM IPTG was added and the culture was incubated for additional 4 hours at 30°C. Harvested cells were sonicated in 50 mM Tris-Cl pH 7.5, 300 mM NaCl, 5% glycerol, 1 mM DTT, 5 mM EDTA, centrifuged 20 min at 20000 g and recombinant GST-tagged proteins were purified using the GST-Bind^TM^ Kit (Novagen). To synthetize the NOVA2 RNA target, pcDNA3 vector containing the sequence CTAGCG<u>TCAT</u>T<u>TCAT</u>C<u>TCAC</u>CA cloned between the *Nhe*1 and *Hin*D3 restriction sites was linearized by *Eco*R1 restriction and 100 ng were transcribed using T7 transcription kit (Ambion) in the presence of 1 µl of [aP^32^]-UTP (Perkin Elmer). It was analyzed on 8% denaturing polyacrylamide and quantified with LS-6500 counter (Beckman). After transcription, 1 unit of DNase I (Invitrogen, Carlsbad, CA) was added, and the sample was incubated for additional 30 min at 37°C. Transcribed RNAs were then purified by micro Bio-Spin 6 chromatography columns (Bio-rad) according to the manufacturer’s instructions. The sizes of RNAs were checked by gel electrophoresis on a denaturing 6% polyacrylamide gel. 10 pM (3000 CPM) of labeled RNA was incubated at 90°C for 5 min in binding buffer (BB, 0.75 mM MgCl2, 50 mM Tris-HCl (pH 7.0), 75 mM NaCl, 37.5 mM KCl, 5.25 mM DTT, 0.1 mg/mL BSA, 0.1 mg/mL Bulk tRNA) and allowed to cool to room temperature. After cooling, RNAsin was added to a final concentration of 0.4 U/μl. Increasing amounts of GST-NOVA2 were then added and the mixture was incubated on ice for 20 min. The solution mixture was loaded onto a non-denaturing 6.0% (w/v) polyacrylamide gel (acrylamide/bisacrylamide, 40:1, w/w) containing 0.5X TBE (1X TBE is 90 mM Tris-base, 89 mM Boric acid and 2 mM EDTA, pH 8.0), which had been pre-electrophoresed at 110 V for 20 min. at 4°C. The gel electrophoresis was run at 110 V at 4°C for 3 hours. Gel was then dried, exposed to a phosphorimager screen, and imaged using a Typhoon 9410. The data were fit to the following equation: y=min+((max-min)/(1+(x/IC50)-HillSlope)) where y is the percentage of RNA bound, x is the concentration of protein, min and max are the minimum and maximum percentage of RNA bound to NOVA2 (0-100%) and IC50 is the concentration where 50% of maximum binding is achieved.

### Zebrafish experiments

Zebrafish (Danio rerio) were raised and maintained as previously described^41^. We injected 6ng of morpholino targeting *nova1* (5’-TCGCCAACTGTCGAGCTTACCTCC-3’) either alone or with 25pg of mRNA, and 50pg of mRNA alone into wildtype zebrafish eggs (AB strain) at the 1-2 cell stage. At 4 days post fertilization, larvae were scored for defects in the number of the inter tecta axonal tracts. Wild-type (ENST00000263257.6) and mutant Mut1 (p.Val261Glyfs*135) full-length human messages were Sanger sequenced and cloned into the pCS2+ vector and transcribed in vitro using the SP6 Message Machine kit (Ambion). We performed a whole mount immunostaining with the anti-acetylated tubulin monoclonal antibody in order to examine the integrity of the inter-tecta axonal tracts of the zebrafish larvae. At 4 days post fertilization, the larvae were fixed in Dent’s solution (80% methanol, 20% dimethylsulphoxide [DMSO]) overnight. After rehydration in decreasing concentration of methanol in PBS, larvae were washed with PBS, permeabilized with 10 μg/mL proteinase K, and postfixed with 4% PFA. Larvae were then washed twice with IF buffer (0.1% Tween-20, 1% BSA in 1× PBS) for 10 min at room temperature. After incubation in blocking solution (10% FBS, 1% BSA in 1× PBS) for 1 hour at room temperature, larvae were incubated with the anti-acetylated tubulin (1:1000) in blocking solution overnight at 4°C. After two washes in IF buffer for 10 min each, larvae were incubated in the secondary antibody solution, 1:500 Alexa Fluor rabbit anti-mouse IgG (Invitrogen), in blocking solution for 1 hour at room temperature and in the dark. After imaging at least 30 larvae per condition from a lateral view on a MacroFluo ORCA Flash (Leica), we scored the number of inter tecta axonal tracts in the brain on each larva for the injected conditions and age-matched controls from the same clutch. All the experiments were repeated 3 times and a t-test was performed to determine significance.

## SUPPLEMENTAL DATA

Supplemental Data include nine figures, two tables and detailed clinical description of the subjects.

## ACKNOWLEDGMENTS

The authors thank the families for their participation to the study. The authors also thank the Fondation Jerome Lejeune, Fondation Maladies Rares and Fondattion APLM for their financial support. This study was also supported by the grant ANR-10-LABX-0030-INRT, a French State fund managed by the Agence Nationale de la Recherche under the frame program Investissements d’Avenir ANR-10-IDEX-0002-02. The authors want also to thank all the people from the IGBMC sequencing platform (Céline Keime, Serge Vicaire, Bernard Jost, Stéphanie Le Gras, Mathieu Jung, etc) and more especially Damien Plassard, for their technical and bioinformatics supports. They thank Jean Muller and Véronique Geoffroy for developing and running Varank on whole-exome sequencing data, as well as Paola Rossadillo and Karim Essabri for their help for the cloning and mutagenesis. They thank Alexandra Benchoua and Istem for providing cells and technical support for hNSCs culture. They also thank people for the Molecular genetic Unit of Strasbourg Hospital (Claire Feger) for Sanger sequencing.

## AUTHOR CONTRIBUTIONS

A.P., F.M. and J-L.M. conceived the study, F.M., B.I., F.T.M-T, N.J., A.T., A.B., C.G., A.G., P.V-N, G.D., Y.C.S., J.Ch., S.G., S.N., S.B., Z.S., E.K., R.D.identified mutations and/or collect patients’ clinical information. F.M., A.P., J.Co., N.D., M-V.H. and A.Q. performed the functional experiments (cell culture, RNAseq, splicing analysis). G.H. and C.G. performed and analyzed zebrafish experiments. C.S. and N.C-B. performed and analyzed immunoflorescence experiments and EMSA. A.P. supervised the study. F.M. and A.P. wrote the manuscript. All authors revised the manuscript.

## WEB RESSOURCES

The URLs for online tools and data presented herein are:

Clinvar: http://www.ncbi.nlm.nih.gov/clinvar/

dbSNP: http://www.ncbi.nlm.nih.gov/projects/SNP/

Decipher: https://decipher.sanger.ac.uk/

ExAC Browser (Beta) | Exome Aggregation Consortium: http://exac.broadinstitute.org/

Exome Variant Server, NHLBI Exome Sequencing Project (ESP): http://evs.gs.washington.edu/EVS/

GeneMatcher: https://genematcher.org/

GEO: https://www.ncbi.nlm.nih.gov/geo/

GnomAD: http://gnomad.broadinstitute.org/

Integrative Genomics Viewer (IGV): http://www.broadinstitute.org/igv/

Mutation Nomenclature: http://www.hgvs.org/mutnomen/recs.html

OMIM: http://www.omim/org/

UCSC: http://genome.ucsc.edu/

Database of Genomic Variants (DGV): http://dgv.tcag.ca/dgv/app/home

