## Supplementaries for "Highly clustered *de novo* frameshift variants in the neuronal splicing factor *NOVA2* result in a specific abnormal C terminal part and cause a severe form of intellectual disability with autistic features"

**Individual 1** was the second child of non-consanguineous parents. The family history was unremarkable. The pregnancy was complicated by hydramnios. He was delivered at term and at birth he had a normal mensuration. He showed neonatal feeding difficulties. He had no dysmorphic features. He was not able to sit independently until 1 year of age and showed truncal hands stereotypic movements. First episode of seizure was noted at the age of 2 years and 6months. He was noted to smile a lot having frequent unprovoked laughter and being attracted to water.  At last evaluation at the age of 9 years and 6 months, he had no speech and could not walk. Physical examination showed lower limbs hypertonia with brisk reflexes. His height was 122cm (-2SD), weight 21kg (-2,5SD) and head circumference 50 cm (-2,5SD). Brain MRI showed cortical atrophy. Angelman syndrome was suspected but 15q11-13 methylation was normal and *UBE3A* direct sequencing did not identify any mutation. Myoclonic epilepsy was treated by sodium valproate.

**Individual 2** was the first child of non-consanguineous parents. The family history was unremarkable. The pregnancy was uncomplicated, with normal screening ultrasounds. He was delivered via vaginal delivery and mensurations at birth were normal. He was hypotonic without facial dysmorphism.  He was not able to sit independently until 1 year of age and walk until 3 years and 6 moths. He smiled a lot and was hyperactive. Physical examination showed brisk lower limbs reflexes and ataxic gait. His height was 105 cm (-2SD), weight 16kg (-2,5SD) and head circumference 49,3 cm (-2,2SD). He showed stereotypic hand movements. MRI of the brain was normal. His behavior and frequent outbursts of anger suggested autism spectrum disorder. Angelman syndrome was suspected but 15q11-13 methylation study was normal.

**Individual 3** was the first the fourth child of non-consanguineous parents. Family history was negative for neurodevelopmental phenotypes. The pregnancy was uncomplicated and patient was born via C-section at 37 weeks gestation. Neonatal course was benign, although sagittal craniosynostosis was not early in infancy. Developmental concerns became apparent to family around 12 months of age, due to speech and gross motor delay. Neurologic evaluation at 2 years of age was notable for hypotonia, Chiari malformation type I, and global developmental delay. Brain MRI at 2 years of age identified stable Chiari I malformation and congenital absence of the left internal carotid artery. The patient had normal female SNP microarray. At her last clinical evaluation at 4 years and 8 months of age, she had been clinically diagnosed with autism spectrum disorder. Physical examination showed microcephaly, ears with pointed helices, ataxia, and hypotonia. She had impaired social interaction with delayed speech for age. Growth parameters were as follows height 103cm (29%), weight 17.7kg (57%). Patient was benefitting from speech, occupational, and music therapy.

**Individual 5** is a 22 year old right handed male with a treatment resistant epilepsy, mental retardation, global developmental delay, and autism. According to his mother, he was born to a 27 year old mother and 33 year old father following an uneventful pregnancy and delivery. His immediate postnatal development seemed normal. However, at about 8 months of age, mother felt that "something was wrong" as he was developing slower as compared to his older brother and cousins. He started walking independently at two years of age, babbled some words but never developed language. At the age of three, he was formally diagnosed with a global developmental delay and autism was diagnosed at nine years of age around the time of the onset of his myoclonic astatic epilepsy. His most disabling seizure types are drop attacks occurring about 15 times a months. His second most common seizure type starts with a fast movement followed by rocking back and forth while remaining semiresponsive. Parents also reported occasional starring spells occurring on average once a month. The patient had a corrective surgery for strabismum at 12 months. He is ambulatory but nonverbal and requires total care and assistance with all activities of daily living, sometimes he would use a spoon for eating. He lives with his family and attends an adult learning center during the daytime. There is no history of ID in the family. A gadolinium enhanced brain MRI performed at 17 years of age showed an absence of neuronal migrational defects. It was remarkable only for a slight cerebellar volume loss, thinning of the an otherwise normally configured corpus callosum in keeping with a global white matter volume loss, most pronounced in biparietal regions. Due to frequent nocturnal awakenings, he was referred for a sleep study at 16 years of age. During the study, interictal discharges in the midline central region were noted but no electrographic or electro-clinical seizures were recorded. There were no central or obstructive apneas. While oxygenation was maintained at or above 96%, significant non-obstructive hypercapnia with pCO2 over 52mmHg was recorded. At the time of the sleep study, his height was 181.3 cm, weight 48.7 kg and the body mass index was 14.82 kg/(m^2). EKG lead documented a normal sinus rhythm throughout the study. He had EEG studies done as a child but the results were not available for review. His general examination was unremarkable. There was no apparent craniofacial dysmorphology and no abnormal cutaneous lesions. His neurological evaluation was remarkable for a poor eye contact and lack of language. He was awake and oriented to his parents, exhibiting behavioral stereotypies manifested by body rocking movements. Cranial nerve and motor examination was unremarkable, deep tendon reflexes were present throughout and pathological reflexes were absent. Sensory evaluation was intact to light touch otherwise not testable. Coordination testing was limited by patient's inability to cooperate but there was no gross ataxia or dysmetria or appendicular tremor, gait was intact. Romberg was negative. Genetic testing performed upon transition to the Adult Epilepsy Clinic included chromosomal microarray and whole exome sequencing including mitochondrial sequence analysis.

**Individual 6** was the second child of non-consanguineous parents. The family history was unremarkable. The pregnancy was uncomplicated. Mensurations at birth were normal. Developmental delay was reported from the age of 6 months, with hypotonia. She was not able to sit independently until 1 year and 3 months and to walk until 3 years and 9 moths. At last evaluation at the age of 5 years and 5 months, she had no speech but could understand simple sentences. Feeding difficulties were reported with hypersensitivity to some textures. She showed mild ataxic gait and flapping hands stereotypic movements. Physical examination showed mild dysmorphic features with deeply set eyes, anteverted nares, and deeply grooved philtrum. Her height was 108 cm (0 SD), weight 16kg (-1SD) and head circumference 48 cm (-2SD). Lower limbs reflexes were normal. MRI of the brain was normal at the age of 9 months. Angelman syndrome was suspected but 15q11-13 methylation study was normal.

**Table S1. List of genes differentially expressed (DE) occurring after *NOVA2* inactivation in hNSCs**

**Table S2. List of the different splicing events occurring after *NOVA2* inactivation in hNSCs, identified by LeafCutter program.**

**Figure S1. Alignement of wild type and the different mutant NOVA2 proteins using ClustalW 2.1.** In grey, the common amino acid sequences introduced by all the frameshift variants. The KH domains (green for KH1, pink for KH2 and blue for KH3) and the position of the Tyr231 are represented. "*" indicates the amino acids common between the mutant and wild-type proteins. Mut1 corresponds to c.782del, p.(Val261Glyfs*135) (identified in Individual 1), Mut2 to c.711_712insTG, p.(Val238Cysfs*159) (identified in Individual 2), Mut3 to c.701_720dup20, p.(Ala241Profs*162) (identified in Individual 3), Mut4 to c.709_748del40, p.(Val237Profs*146) (identified in Individual 4), Mut5 to c.781del, p.(Val261Trpfs*135) (identified in Individual 5), Mut6 to c.720_721insCCGCGGATGTGCTTCCAGCC, p.(Ala241Profs*162)(identified in Individual 6).

**Figure S2. Predicted structures for wild-type and mutant NOVA2 protein**

The 3D structures were generated from FASTA protein sequences using RaptorX for NOVA2 wild-type (WT), and mutant NOVA2 protein carrying patient's variant Mut1 or carrying a stop codon Tyr231*. Predicted alpha-helices and beta-sheets are represented. The three KH domains are indicated (green for KH1, pink for KH2 and blue for KH3).

**Figure S3. Expression and cellular localization of Mut1 NOVA2 proteins in HeLa cells**

(**a**) Expression of NOVA2 in HeLa cells. HeLa cells were co-transfected with EGFP-tagged NOVA2 wild-type (WT) or mutant cDNA and a plasmid with a FLAG-tagged protein as a control for transfection. Cells were harvested 24 h after transfection, and expression of NOVA2 was analyzed by SDS–PAGE and immunoblotting with anti-GFP and anti-FLAG antibodies. Quantification of NOVA2 expression: the ratio between GFP and the transfection control FLAG was calculated from four independent experiments and each condition was compared to the WT. The error bars indicate the standard error mean (SEM). Kruskal-Wallis ANOVA with Dunn's multiple comparisons * $p<0.05$, ns: non-significant (**b)** Cellular localization of NOVA2 proteins in Hela cells. Hela cell lines were transiently transfected with HA-NOVA2 WT or HA-NOVA2 Mut1 and immunofluorescence experiments using an anti-HA antibody revealed a nuclear localization of both WT and mutant NOVA2 proteins. The DAPI (4′,6- diamidino- 2- phenylindole) staining indicates the position of the nuclei.

**Figure S4. Differential expression after *NOVA2* inactivation in hNSCs**

The graph on the left represents a scatterplot of the normalized gene expression. For a given gene, its value is obtained by mean on all samples belonging to the same condition. The graph on the right represents a volcano plot. The x-axis represents the log2 Fold Change values and the y-axis the p-values (in -log10). Red points show values with an adjusted p-value < 0.05 and orange points show values with an adjusted p-value < 0.1. Expression in (**a)** hNSCs treated with non-specific scramble siRNA compared to untreated cells (INT) and expression in (**b)** hNSCs treated with *NOVA2* scramble siRNA compared to untreated cells.

**Figure S5, S6 and S7. Additional examples of alternative splicing events regulated by NOVA2 in human cells*:* *NEO1* exon 26 (S5)*,* *SORBS1* exon 3 (S6) and *AKAP13* exon 12 (S7).**

(**a**) Sashimi plot established from RNA-Seq data for *NEO1* exon 26, *SORBS1* exon 3 and *AKAP13* exon 12. Sashimi plot established from RNA-Seq data are represented. The number of reads supporting the existence of each exon-exon junction is indicated as an average between data from the two independent series of hNSCs treated with Interferin only (in red), with scramble siRNA (in blue) or with *NOVA2* siRNA (in green) during 48 hours. **(b)** Confirmation of the consequences of *NOVA2* inactivation on these AS events in another hNSC cell line (GM01869), transfected with siScramble (si Scr), si*NOVA2*, or treated with the transfecting agent only (INT). The RT-PCR products obtained were analyzed by migration on a 2,100 Bioanalyzer instrument (Agilent Technology). Experiments were done in triplicates The error bars indicate the SEM. Kruskal–Wallis' ANOVA with Dunn's multiple comparison testwas performed * p<0.05, ns: non-significant. **(c)** Effect of overexpression of NOVA2 proteins on these AS events in HeLa cells transfected with EGFP-tagged NOVA2 wild-type (WT) or mutant cDNAs. The RT-PCR products obtained were analyzed by migration on a 2,100 Bioanalyzer instrument (Agilent Technology). Four series of experiments were analyzed. The error bars indicate the SEM. Brown-Forsythe and Welch's ANOVA with Holm-Sidak's multiple comparisons *** p<0.001, **p<0.01, * p<0.05, ns: non-significant.

**Figure S8. Effect of co-expression of wild-type and mutant NOVA2 proteins on the regulation of AS events in HeLa cells**

HeLa cells were transfected with empty plasmid, EGFP-tagged NOVA2 wild-type and mutants proteins alone or combined together. The RT-PCR products obtained were analyzed by migration on a 2,100 Bioanalyzer instrument (Agilent Technology). Three series of experiments were analyzed. The error bars indicate the SEM. Kruskal–Wallis' ANOVA with Dunn's multiple comparison test was performed ns: non-significant.

**Figure S9. Depth of coverage along the *NOVA2* gene obtained by WES (extracted from gnomAD)** Representation of the depth of coverage of *NOVA2* coding sequences which can be obtained by Whole Exome Sequencing (WES, in blue) or Whole Genome Sequencing (WGS, in green)(data extracted from gnomAD: https://gnomad.broadinstitute.org/gene/ENSG00000104967). Two regions of exon 4 are poorly (<10X, red line) or not covered: c.720 to 750, encoding amino acids 240 to 250, and c.1005 to 1140 encoding amino acids 355 to 380.

```
NOVA2     MEPEAPDSRKRPLETPPEVVCTKRSNTGEEGEYFLKVLIPSYAAGSIIGKGGQTIVQLQK      60

Mut1      MEPEAPDSRKRPLETPPEVVCTKRSNTGEEGEYFLKVLIPSYAAGSIIGKGGQTIVQLQK      60
Mut2      MEPEAPDSRKRPLETPPEVVCTKRSNTGEEGEYFLKVLIPSYAAGSIIGKGGQTIVQLQK      60
Mut3      MEPEAPDSRKRPLETPPEVVCTKRSNTGEEGEYFLKVLIPSYAAGSIIGKGGQTIVQLQK  KH1 60
Mut4      MEPEAPDSRKRPLETPPEVVCTKRSNTGEEGEYFLKVLIPSYAAGSIIGKGGQTIVQLQK      60
Mut5      MEPEAPDSRKRPLETPPEVVCTKRSNTGEEGEYFLKVLIPSYAAGSIIGKGGQTIVQLQK      60
Mut6      MEPEAPDSRKRPLETPPEVVCTKRSNTGEEGEYFLKVLIPSYAAGSIIGKGGQTIVQLQK      60
          ************************************************************

NOVA2     ETGATIKLSKSKDFYPGTTERVCLVQGTAEALNAVHSFIAEKVREIPQAMTKPEVVNILQ      120

Mut1      ETGATIKLSKSKDFYPGTTERVCLVQGTAEALNAVHSFIAEKVREIPQAMTKPEVVNILQ      120
Mut2      ETGATIKLSKSKDFYPGTTERVCLVQGTAEALNAVHSFIAEKVREIPQAMTKPEVVNILQ      120
Mut3      ETGATIKLSKSKDFYPGTTERVCLVQGTAEALNAVHSFIAEKVREIPQAMTKPEVVNILQ      120
Mut4      ETGATIKLSKSKDFYPGTTERVCLVQGTAEALNAVHSFIAEKVREIPQAMTKPEVVNILQ      120
Mut5      ETGATIKLSKSKDFYPGTTERVCLVQGTAEALNAVHSFIAEKVREIPQAMTKPEVVNILQ      120
Mut6      ETGATIKLSKSKDFYPGTTERVCLVQGTAEALNAVHSFIAEKVREIPQAMTKPEVVNILQ      120
          ************************************************************

NOVA2     PQTTMNPDRAKQAKLIVPNSTAGLIIGKGGATVKAVMEQSGAWVQLSQKPEGINLQERVV      180

Mut1      PQTTMNPDRAKQAKLIVPNSTAGLIIGKGGATVKAVMEQSGAWVQLSQKPEGINLQERVV      180
Mut2      PQTTMNPDRAKQAKLIVPNSTAGLIIGKGGATVKAVMEQSGAWVQLSQKPEGINLQERVV      180
Mut3      PQTTMNPDRAKQAKLIVPNSTAGLIIGKGGATVKAVMEQSGAWVQLSQKPEGINLQERVV  KH2 180
Mut4      PQTTMNPDRAKQAKLIVPNSTAGLIIGKGGATVKAVMEQSGAWVQLSQKPEGINLQERVV      180
Mut5      PQTTMNPDRAKQAKLIVPNSTAGLIIGKGGATVKAVMEQSGAWVQLSQKPEGINLQERVV      180
Mut6      PQTTMNPDRAKQAKLIVPNSTAGLIIGKGGATVKAVMEQSGAWVQLSQKPEGINLQERVV      180
          ************************************************************

                                                     Y231
NOVA2     TVSGEPEQVHKAVSAIVQKVQEDPQSSSCLNISYANVAGPVANSNPTGSPYASPADVLPA      240

Mut1      TVSGEPEQVHKAVSAIVQKVQEDPQSSSCLNISYANVAGPVANSNPTGSPYASPADVLPA      240
Mut2      TVSGEPEQVHKAVSAIVQKVQEDPQSSSCLNISYANVAGPVANSNPTGSPYASPAD----      236
Mut3      TVSGEPEQVHKAVSAIVQKVQEDPQSSSCLNISYANVAGPVANSNPTGSPYASPADVLPA      240
Mut4      TVSGEPEQVHKAVSAIVQKVQEDPQSSSCLNISYANVAGPVANSNPTGSPYASPAD----      236
Mut5      TVSGEPEQVHKAVSAIVQKVQEDPQSSSCLNISYANVAGPVANSNPTGSPYASPADVLPA      240
Mut6      TVSGEPEQVHKAVSAIVQKVQEDPQSSSCLNISYANVAGPVANSNPTGSPYASPADVLPA      240
          ********************************************************

NOVA2     AAAASAAAASGLLGPAGLAGVGAFPAALPAFSGTDLLAISTALNTLASYGYNTNSLGLGL      300

Mut1      AAAASAA-------AASGLLGPAGLAGGGPFPPRCPPSQAPTCWPSARRLTRWQVTATTP      293
Mut2      --VCCQPRPQRRPPPPPACWAPPGWLAWGPFPPRCPPSQAPTCWPSARRLTRWQVTATTP      294
Mut3      PRMCCQPRPQRRPPPPPACWAPPGWLAWGPFPPRCPPSQAPTCWPSARRLTRWQVTATTP      300
Mut4      ----------------PACWAPPGWLAWGPFPPRCPPSQAPTCWPSARRLTRWQVTATTP      280
Mut5      AAAASAA-------AASGLLGPAGLAGWGPFPPRCPPSQAPTCWPSARRLTRWQVTATTP      293
Mut6      PRMCFQPRPQRRPPPPPACWAPPGWLAWGPFPPRCPPSQAPTCWPSARRLTRWQVTATTP      300

NOVA2     NSAAASGVLAAVAAGANPAAAAAANLLASYAGEAGAGPAGGAAPPPPPPPGALGSFALAA      360

Mut1      TPWAWASTRPQLPASWPPWPPGPTQQPPPPPTSWHPTRARPGPGQPEGPPRRRPRLPEPW      353
Mut2      TPWAWASTRPQLPASWPPWPPGPTQQPPPPPTSWHPTRARPGPGQPEGPPRRRPRLPEPW      354
Mut3      TPWAWASTRPQLPASWPPWPPGPTQQPPPPPTSWHPTRARPGPGQPEGPPRRRPRLPEPW      360
Mut4      TPWAWASTRPQLPASWPPWPPGPTQQPPPPPTSWHPTRARPGPGQPEGPPRRRPRLPEPW      340
Mut5      TPWAWASTRPQLPASWPPWPPGPTQQPPPPPTSWHPTRARPGPGQPEGPPRRRPRLPEPW      353
Mut6      TPWAWASTRPQLPASWPPWPPGPTQQPPPPPTSWHPTRARPGPGQPEGPPRRRPRLPEPW      360

NOVA2     AANGYLGAGAGGGAGGGGGPLVAAAAAAGAAGGFLTAEKLAAESAKELVEIAVPENLVGA  KH3 401

Mut1      GPLRWPQPPTATSGPGRAAGRAEGAARWWPLQPRPGRPGAS   394
Mut2      GPLRWPQPPTATSGPGRAAGRAEGAARWWPLQPRPGRPGAS   395
Mut3      GPLRWPQPPTATSGPGRAAGRAEGAARWWPLQPRPGRPGAS   401
Mut4      GPLRWPQPPTATSGPGRAAGRAEGAARWWPLQPRPGRPGAS   381
Mut5      GPLRWPQPPTATSGPGRAAGRAEGAARWWPLQPRPGRPGAS   394
Mut6      GPLRWPQPPTATSGPGRAAGRAEGAARWWPLQPRPGRPGAS   401

NOVA2     ILGKGGKTLVEYQELTGARIQISKKGEFLPGTRNRRVTITGSPAATQAAQYLISQRVTYE      480

NOVA2     QGVRASNPQKVG       492
```

**Figure S1**

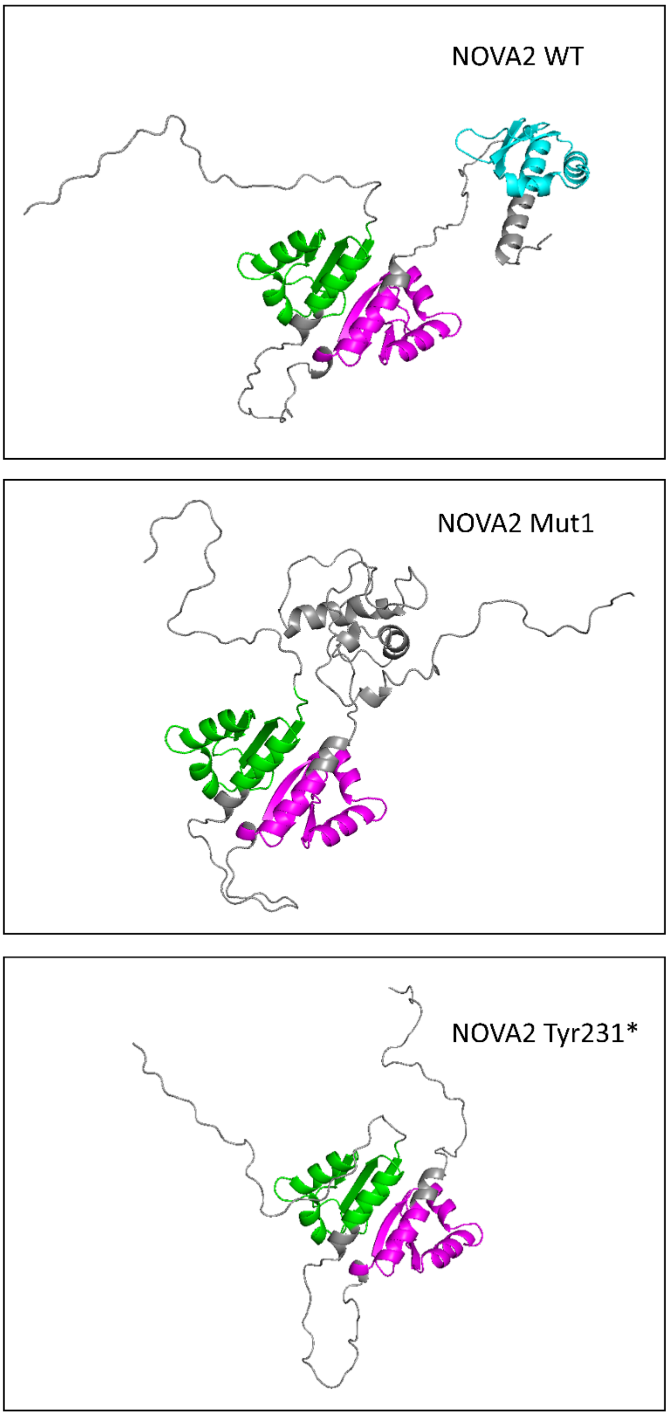


**Figure S2**

a

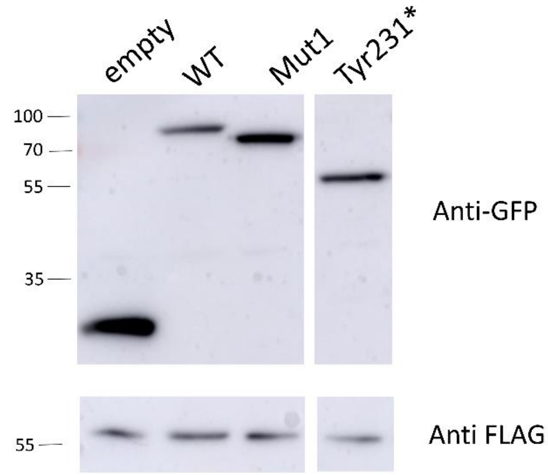
empty
WT
Mut1
Tyr231*
100
70
55
35
Anti-GFP
55
Anti FLAG

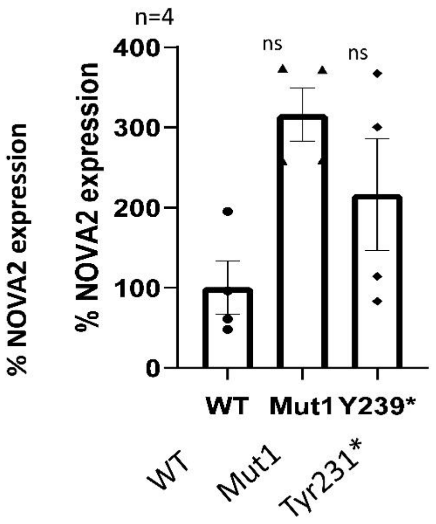
n=4
400
300
200
100
0
ns
ns
% NOVA2 expression
% NOVA2 expression
WT
Mut1
Y239*
WT
Mut1
Tyr231*

b

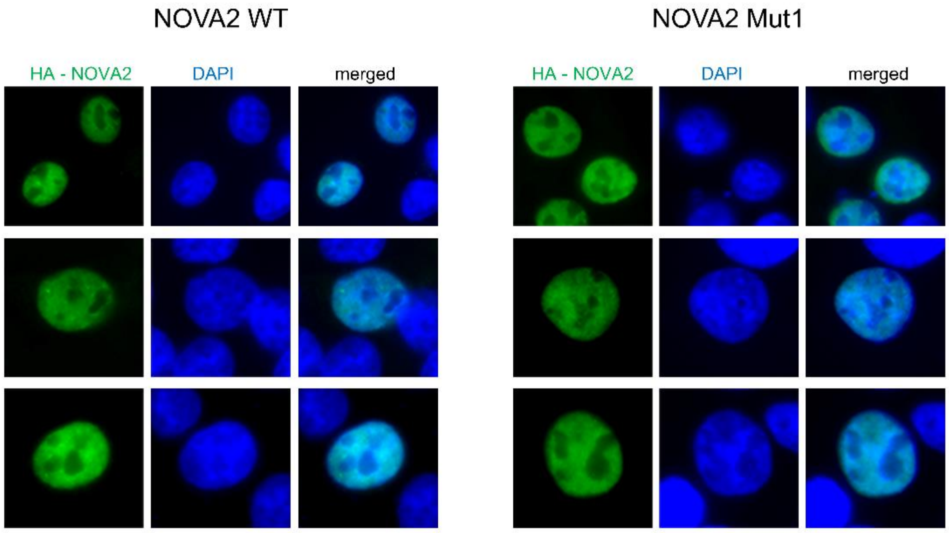
NOVA2 WT
NOVA2 Mut1
HA - NOVA2
DAPI
merged
HA - NOVA2
DAPI
merged

**Figure S3**

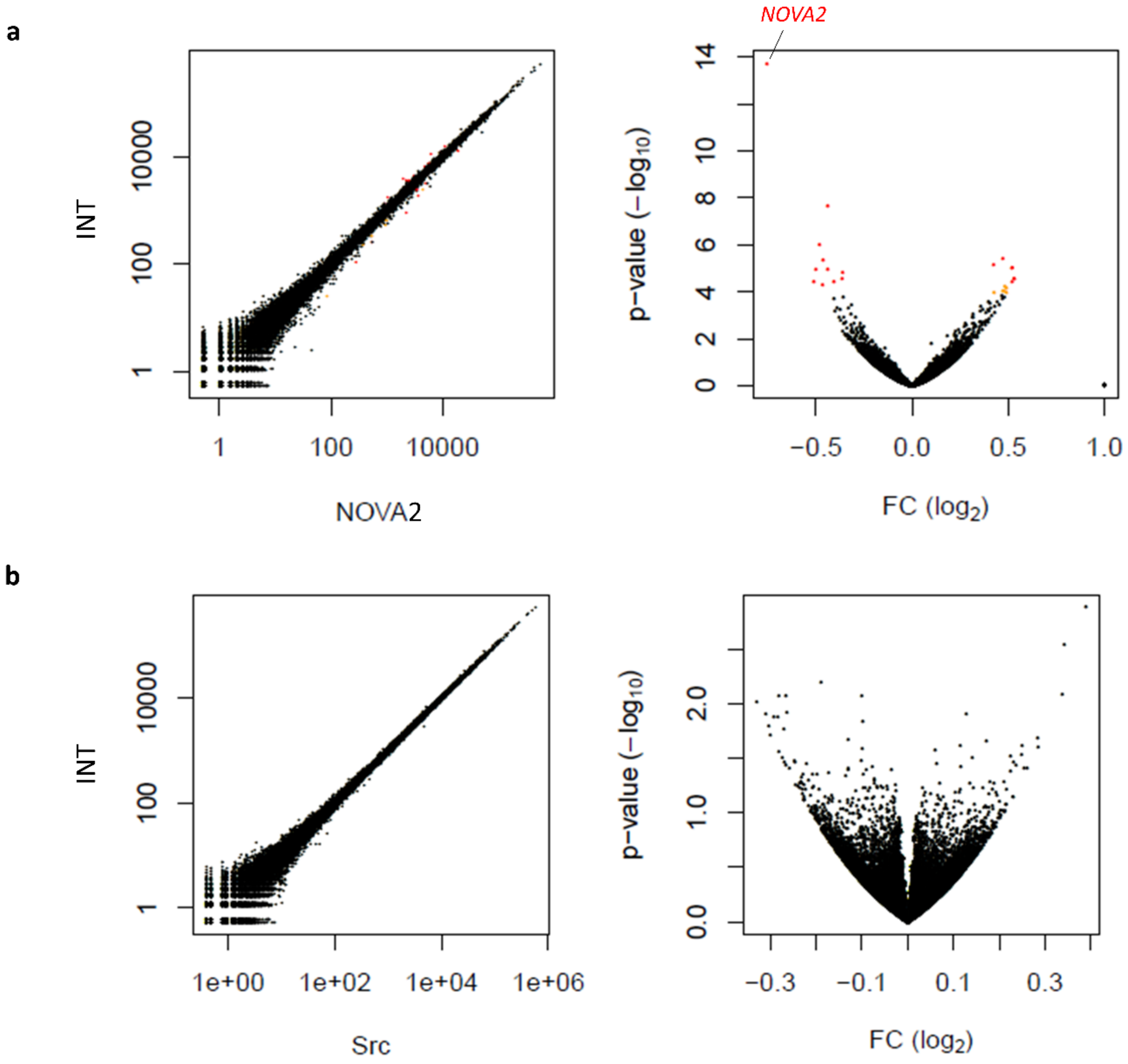


**Figure S4**

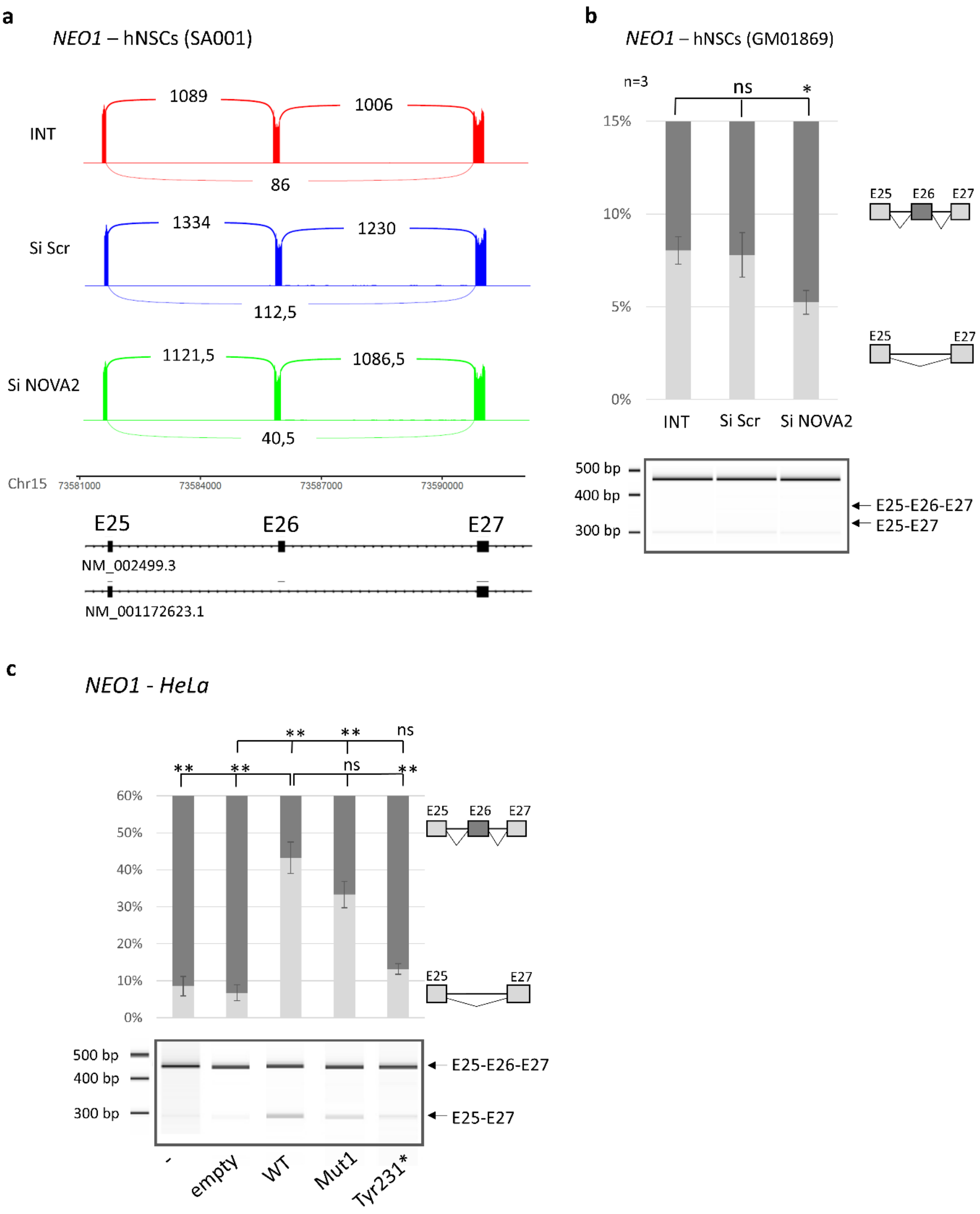
a
NEO1 – hNSCs (SA001)
INT
1089
1006
86
Si Scr
1334
1230
112,5
Si NOVA2
1121,5
1086,5
40,5
Chr15
73581000
73584000
73587000
73590000
E25
E26
E27
NM_002499.3
NM_001172623.1
b
NEO1 – hNSCs (GM01869)
n=3
ns
*
15%
10%
5%
0%
INT
Si Scr
Si NOVA2
E25
E26
E27
500 bp
400 bp
300 bp
E25-E26-E27
E25-E27
c
NEO1 - HeLa
**
ns
60%
50%
40%
30%
20%
10%
0%
-
empty
WT
Mut1
Tyr231*

**Figure S5**

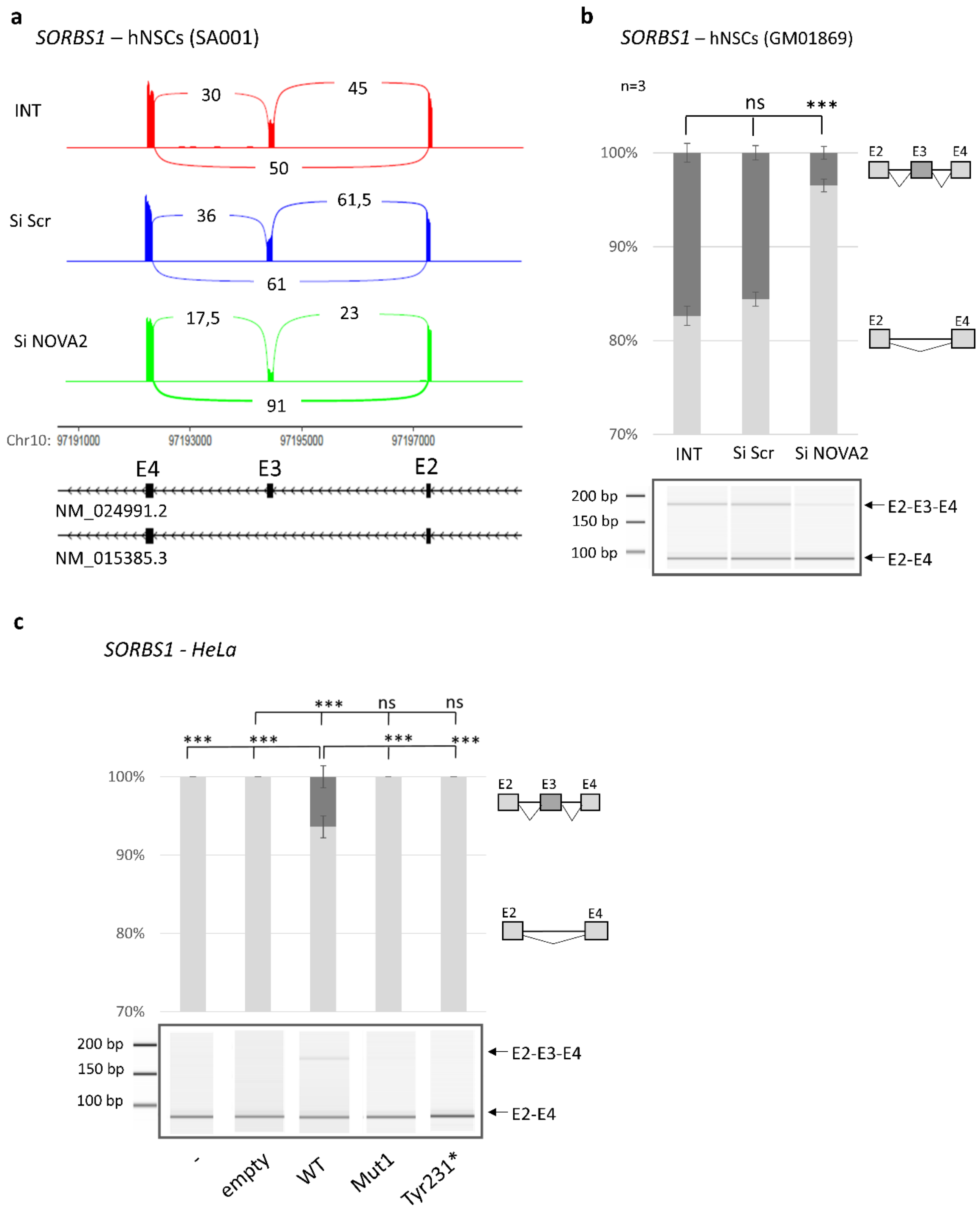
a
SORBS1 – hNSCs (SA001)
INT
30
45
50
Si Scr
36
61,5
61
Si NOVA2
17,5
23
91
Chr10: 97191000
97193000
97195000
97197000
E4
E3
E2
NM_024991.2
NM_015385.3
b
SORBS1 – hNSCs (GM01869)
n=3
ns
***
100%
90%
80%
70%
E2
E3
E4
INT
Si Scr
Si NOVA2
200 bp
150 bp
100 bp
E2-E3-E4
E2-E4
c
SORBS1 - HeLa
***
ns
100%
90%
80%
70%
-
empty
WT
Mut1
Tyr231*

**Figure S6**

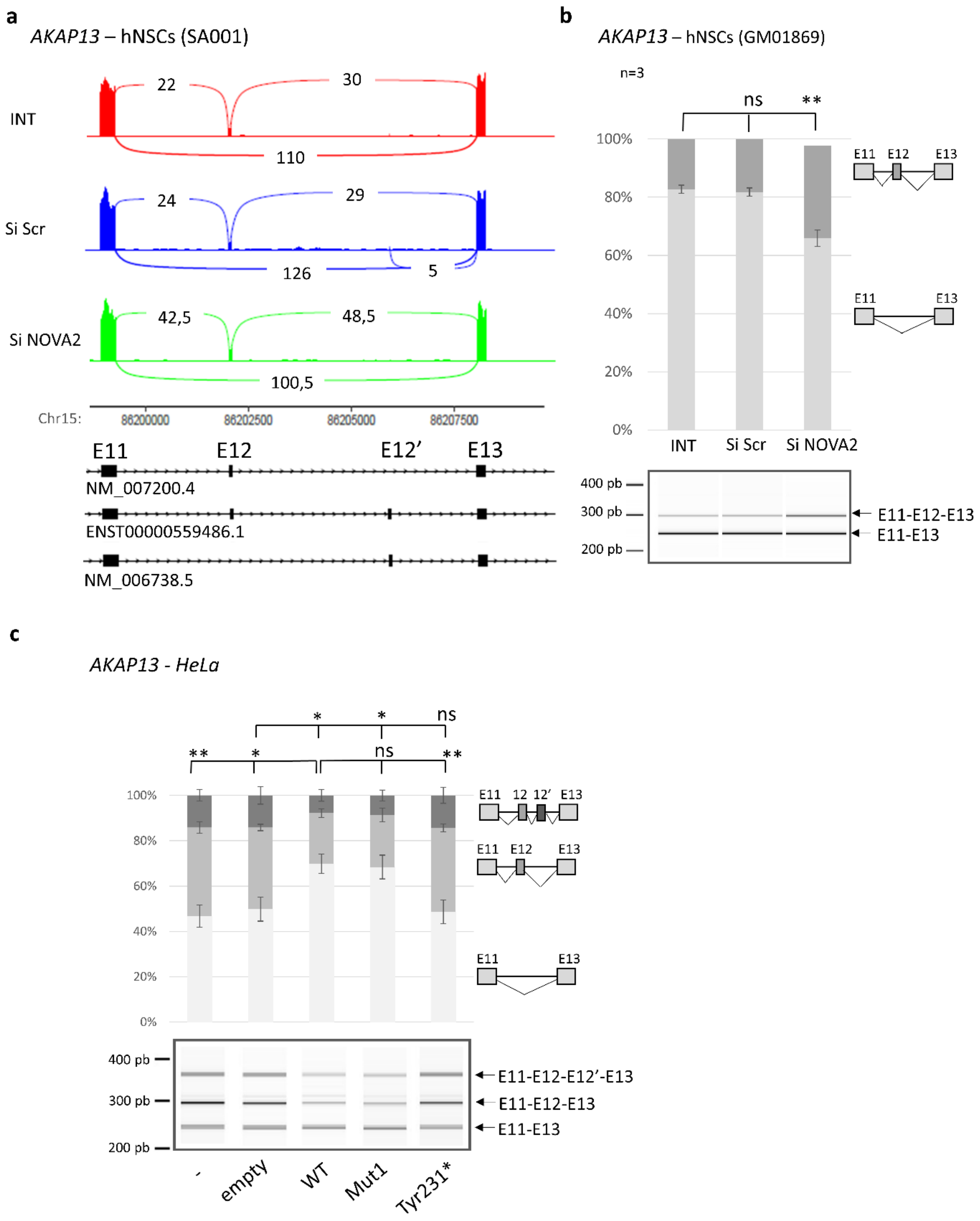
a
AKAP13 – hNSCs (SA001)
INT
Si Scr
Si NOVA2
22
30
110
24
29
126
5
42,5
48,5
100,5
Chr15:
86200000
86202500
86205000
86207500
E11
E12
E12'
E13
NM_007200.4
ENST00000559486.1
NM_006738.5
b
AKAP13 – hNSCs (GM01869)
n=3
ns
**
100%
80%
60%
40%
20%
0%
INT
Si Scr
Si NOVA2
E11 E12 E13
E11 E13
400 pb
300 pb
200 pb
E11-E12-E13
E11-E13
c
AKAP13 - HeLa
*
*
ns
**
*
ns
**
100%
80%
60%
40%
20%
0%
E11 12 12' E13
E11 E12 E13
E11 E13
400 pb
300 pb
200 pb
E11-E12-E12'-E13
E11-E12-E13
E11-E13
-
empty
WT
Mut1
Tyr231*

**Figure S7**

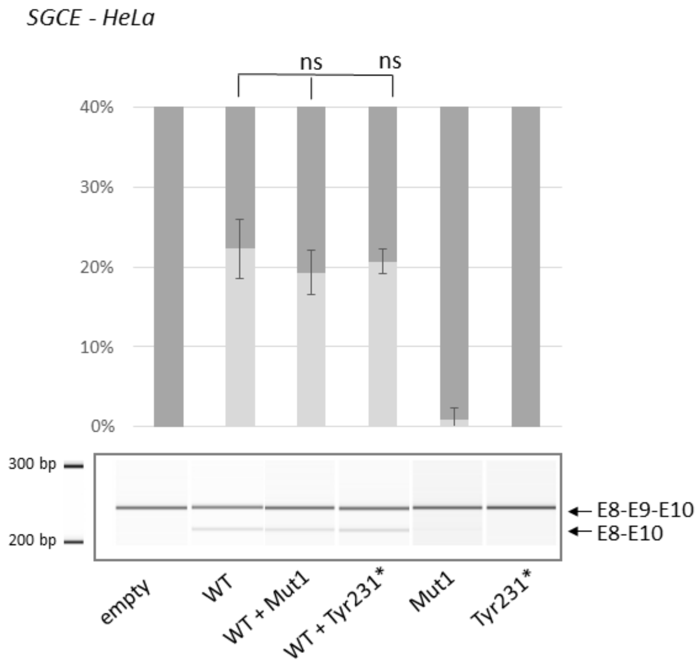
SGCE - HeLa
ns
ns
40%
30%
20%
10%
0%
300 bp
200 bp
E8-E9-E10
E8-E10
empty
WT
WT + Mut1
WT + Tyr231*
Mut1
Tyr231*

**Figure S8**

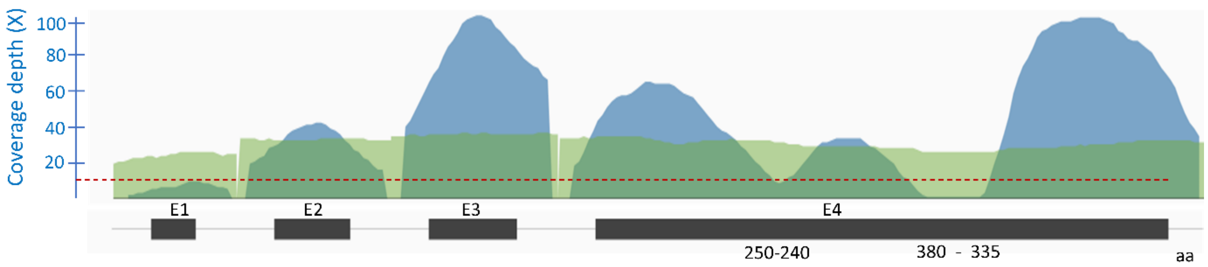


**Figure S9**
